# Catching the big picture of the Mediterranean Sea biodiversity with an end-to-end model of climate and fishing impacts

**DOI:** 10.1101/593822

**Authors:** Fabien Moullec, Laure Velez, Philippe Verley, Nicolas Barrier, Caroline Ulses, Pierluigi Carbonara, Antonio Esteban, Cristina Follesa, Michele Gristina, Angélique Jadaud, Alessandro Ligas, Eduardo López Díaz, Porzia Maiorano, Panagiota Peristeraki, Maria Teresa Spedicato, Ioannis Thasitis, Maria Valls, François Guilhaumon, Yunne-Jai Shin

## Abstract

The Mediterranean Sea is among the main hotspots of marine biodiversity in the world. Under combined pressures of fishing activities and climate change it has also become a hotspot of global change, with increased concern about the worsening status of marine exploited species. More integrated modelling approaches are needed to anticipate global change impacts in the Mediterranean Sea, in order to help decision makers prioritizing management actions and strategies, mitigating impacts and adapting to changes. Our challenge was to develop a holistic model of the marine biodiversity in the Mediterranean Sea with an explicit representation of the spatial multispecies dynamics of exploited resources under the combined influence of climate variability and fishing pressure. An individual-based ecosystem model OSMOSE (Object-oriented Simulator of Marine ecOSystEms), including 100 marine species (fish, cephalopods and crustaceans) and representing about 95 % of the total declared catches, has been implemented for the first time at a high spatial resolution (400 km^2^) and at a large spatial scale (whole Mediterranean basin). The coupling of OSMOSE to the NEMOMED 12 physical model, and to the Eco3M-S biogeochemical and low trophic level model has been achieved to build the OSMOSE-MED end-to-end model. We fitted OSMOSE-MED to observed and estimated data of biomass and commercial catches using a likelihood approach and an evolutionary optimization algorithm. The outputs of OSMOSE-MED were then verified against observed biomass and catches, and confronted to independent datasets (MEDITS data, diet compositions and trophic levels). Although some improvements are suggested for future developments, the model results at different hierarchical levels, from individuals up to the ecosystem scale, were consistent with current knowledge and observations on the structure, the functioning and the dynamics of the ecosystems in the Mediterranean Sea. All the modelling steps, from the comprehensive representation of key ecological processes and feedbacks, the careful parameterization of the model, the confrontation to observed data, and the positive outcome from the validation process, allowed to strengthen the degree of realism of OSMOSE-MED and its relevance as an impact model to explore the futures of marine biodiversity under scenarios of global change, and as a tool to support the implementation of ecosystem-based fisheries management in the Mediterranean Sea.

## 1 Introduction

The Mediterranean Sea is the largest of the semi-enclosed European seas and one of the main reservoirs of biodiversity in the world (Coll et al., 2010). It embeds 4 to 18% of identified marine species which is considerable given it only covers 0.82% of the global ocean surface (Coll et al., 2010). The Mediterranean Sea is also a hotspot of global changes (Coll et al., 2012, 2010; Giorgi, 2006; Giorgi and Lionello, 2008; Micheli et al., 2013a; Ramírez et al., 2018; Stock et al., 2018). Overfishing, pollutions from land-based sources, degradation or loss of critical habitats, species introductions and climate change are all pervasive in the Mediterranean Sea and may operate in synergy, leading to deep modifications of the structure, stability and functioning of marine Mediterranean ecosystems (Albouy et al., 2014; Coll et al., 2012; Lotze et al., 2006). Fishing is probably one of the highest threat for biodiversity in the region: Exploitation rate has been steadily and steeply increasing with poor fishing selectivity, and fish stocks have been shrinking (Colloca et al., 2017; Vasilakopoulos et al., 2014). As a consequence, more than 90 % of the assessed stocks were categorized as overfished in 2017 (GFCM, 2017a; STECF, 2017). Nonetheless, while fish stocks are declining on the continental shelf, especially those of long-living species such as European hake (*Merluccius merluccius*), a few short-living species such as shrimps, cephalopods, and other fish species (e.g., red mullet, *Mullus barbatus*), have shown increasing trends in biomass (GFCM, 2017a; Maynou et al., 2011). Deep-water rose shrimp, *Parapenaeus longirostris*, is the most emblematic example: its biomass has increased all over the Mediterranean Sea in the last decade due to increasing temperature and decrease of predatory pressure (i.e. by European hake) (Colloca et al., 2014; Ligas et al., 2011; Sbrana et al., in press).

In the absence (or lack) of strong management plans, the deteriorating status of fisheries and their resources in the Mediterranean Sea is likely to aggravate, especially in a climate change context (Cheung et al., 2018; FAO, 2018). The Mediterranean Sea has been identified as one of the most vulnerable regions in future climate change projections (Cramer et al., 2018; Giorgi, 2006; Hoegh-Guldberg et al., 2014). Effects of climate change on marine ecosystems are already clearly perceivable, with impacts reported from low (e.g. macrophytes, phytoplankton) to high (e.g. predatory fish) trophic levels, from individual up to the ecosystem scale (Calvo et al., 2011; Durrieu de Madron et al., 2011; Lejeusne et al., 2010; Marbà et al., 2015; Tzanatos et al., 2014) which could affect biodiversity, commercial fisheries, food web and ecosystem functioning (Albouy et al., 2014; AllEnvi, 2016; Bosello et al., 2015; Hattab et al., 2014; Jordà et al., 2012; Marbà et al., 2015; Pecl et al., 2017; Piroddi et al., 2017).

Anthropogenic pressures on Mediterranean ecosystems are projected to increase in the future, especially those related to climate change, habitat degradation and exploitation (Butchart et al., 2010; Calvo et al., 2011; Coll et al., 2010). Considering the diversity of human and natural pressures and the possibility that they act in synergy on marine ecosystems, there is an urgent need for scientists and decision-makers to develop more holistic and integrative approaches to quantify, anticipate, mitigate and manage human impacts on natural environments (Colloca et al., 2017; Hilborn, 2011; Link, 2010). Along these lines, Ecosystem-Based Management (EBM) and more specifically the Ecosystem Approach to Fisheries Management (EAFM) emerged in the early 1990s to consider all anthropogenic activities which could affect the sustainability of goods and services provided by ecosystems (Pikitch et al., 2004). In the European Union seas, these approaches are mainly framed by the Common Fisheries Policy (CFP, 2013), and the European Marine Strategy Framework Directive (MSFD; European Commission, 2008) that requires that all member states take the necessary measures to achieve or maintain Good Environmental Status for marine ecosystems, with the explicit regulatory objective that “biodiversity is maintained” by 2020 at the latest (European Commission, 2008). The MSFD thereby requires the development of suitable tools to evaluate the status of marine ecosystems and their responses to human activities, and to manage and harvest sustainably all commercial species. In this regard, it is essential to develop our capacities in projecting the future impacts of a variety of policy interventions and strategic management plans for restoring marine ecosystems and biodiversity while ensuring sustained provision of marine fisheries products to human societies.

In order to propose plausible biodiversity scenarios at the scale of the whole Mediterranean Sea, which would relevantly support management decision in the region, the challenge that is addressed in this paper is to develop a model able to represent in an explicit way the spatial multispecies dynamics of marine resources under the combined influence of climate change and fishing pressure. The most recent End-to-End models (E2E), representing the entire food web, from plankton to top predators as well as their associated abiotic environment, are expected to provide valuable tools for assessing the effects of climate and fishing on ecosystem dynamics (Fulton, 2010; Grimm et al., 2017; Nicholson et al., 2018; Piroddi et al., 2017, 2015b; Rose et al., 2010; Travers et al., 2007). Notwithstanding the state-of-the-art modelling of food webs and multispecies communities in Mediterranean ecosystems, there is still a gap in modelling the dynamics of biodiversity at the scale of the whole Mediterranean Sea, accounting for the complexity of species introductions, multispecies interactions and spatial dynamics in a global change context. While trophic modelling has improved greatly on coastal marine ecosystems in different parts of the Mediterranean Sea, no study has yet succeeded to model species assemblages at the whole Mediterranean scale with an explicit modelling of the multispecies, spatial, trait-based, whole life cycle dynamics and interactions of a hundred exploited species.

In this paper, we present the individual-based, ecosystem model OSMOSE (Object-oriented Simulator of Marine ecOSystEms) that was used for the first time at large spatial scale (the whole Mediterranean basin), with a high spatial resolution (400 km^2^), and for as many as 100 marine species (fish, cephalopods and crustaceans) representing about 95 % of total declared catches in the Mediterranean Sea. We built an end-to-end modelling approach of the Mediterranean Sea by coupling the OSMOSE model (representing the higher trophic level species) to the physical model NEMOMED 12, and to the biogeochemical model Eco3M-S (representing the low trophic levels). The resulting end-to-end model OSMOSE-MED was calibrated to represent the Mediterranean Sea during the 2006-2013 period. Here, we: (i) start with a brief description of the NEMOMED 12, Eco3M-S and OSMOSE component models; (ii) detail the parameterization of OSMOSE-MED; (iii) present the methodology implemented to calibrate OSMOSE-MED; (iv) evaluate the capacity of OSMOSE-MED to represent some key indicators of the Mediterranean Sea, namely biomass, catches, trophic levels, at the individual up to the community scales; (v) discuss the challenges incurred by the development of such complex end-to-end models as well as associated limitations.

## 2 Materials and Methods

The individual-based model OSMOSE considers a large proportion of the fishable food web and simulates trophic interactions between several target and non-target marine species, mainly fish species. In order to model the effects of environmental heterogeneity and variability which could affect the entire food web by bottom-up control, OSMOSE has been forced (i.e. one way coupling – offline) by the Low Trophic Levels (LTL) NEMOMED 12 / Eco3M-S model. The end-to-end OSMOSE-MED model thus formed represents the whole food web from primary and secondary producers to main top predators.

### 2.1 The low trophic levels model NEMOMED 12 / Eco3M-S

Eco3M-S is a biogeochemical model that simulates the lower trophic levels of marine ecosystems (phyto-and zoo-plankton), the biogeochemical cycles of carbon and other key elements such as phosphorus and nitrogen in the Mediterranean Sea (Auger et al., 2011; Ulses et al., 2016). Independently from our study, Eco3M-S has been coupled to NEMOMED12, a high resolution (≈1/12°) hydro-dynamical model adapted to the Mediterranean region (see Beuvier et al., 2012 for more details on the structure and parameterization of NEMOMED 12) (Kessouri, 2015; Kessouri et al., 2017).

NEMOMED12 is a regional circulation model, which is an updated version of the OPAMED 8 and NEMOMED 8 models used by Ben Rais Lasram et al. (2010), Hattab et al. (2014), Albouy et al. (2014, 2013, 2012) and more recently by Halouani et al. (2016) as input for niche/habitat models at local or regional scale in the Mediterranean Sea. The NEMOMED 12 domain covers the whole Mediterranean Sea and part of the Atlantic Ocean (from 11 °W to 7.5 °W) to take into account the inter-oceans exchanges (Beuvier et al., 2012a; Beuvier et al., 2012b). It does not cover the Black Sea. Based on the standard three-polar ORCA grid of NEMO at 1/12° (≈7 km), NEMOMED 12 resolution varies in latitude and longitude but allows to explicitly resolve most of the mesoscale features. NEMOMED 12 is thus an eddy-resolving model in the major part of the Mediterranean Sea (Beuvier et al., 2012a). It has a time step of 12 minutes, and is daily forced by the atmospheric ARPERA data, obtained by performing a dynamical downscaling of ECMWF (European Centre for Medium-Range Weather Forecasts) products above the European-Mediterranean region (Beuvier et al., 2012a; Herrmann and Somot, 2008).

The coupling between NEMOMED 12 and the biogeochemical Eco3M-S model was done offline (one way coupling). The Eco3M-S model represents several elements’ cycles such as carbon (C), nitrogen (N), phosphorus (P) and silica (Si) in order to reproduce the different limitations and co-limitations observed in the Mediterranean Sea and the dynamics of different plankton groups. Seven planktonic functional types (PFTs) representing the main PFTs and the range of the plankton size spectrum of the Mediterranean Sea were modelled. Thus, the structure of the trophic web base includes three size-classes of phytoplankton (pico-, nano-, and micro-phytoplankton), three size-classes of zooplankton (nano-, micro-, and meso-zooplankton), and heterotrophic bacteria as decomposers (Table 1). The representation of the phytoplankton dynamics was derived from the Eco3M model presented in Baklouti et al. (2006). Among primary producers, nanophytoplankton dominated the biomass of phytoplankton communities for most of the year, and microphytoplankton could punctually contribute to a large part of primary production during the spring period in the Northwestern Mediterranean Sea (Auger et al., 2011; Ulses et al., 2016). The structure of Eco3M-S reflects major grazing links such as nanozooplankton preying on the small phytoplankton group and bacteria, microzooplankton consuming microphytoplankton, and mesozooplankton, mainly composed by copepods, grazing on the largest categories of plankton (i.e. microphyto-and microzoo-plankton). Bacteria (i.e. picoheterotroph plankton) are responsible for the remineralization of the dissolved organic matter. The representation of the heterotrophic processes is based on the models developed by Anderson and Pondaven (2003) and Raick et al. (2005). All features, formulations and parameterization of biogeochemical processes integrated in the mechanistic Eco3M-S model were described in details by Auger et al. (2011), Kessouri (2015) and Ulses et al. (2016).

**Table 1.**
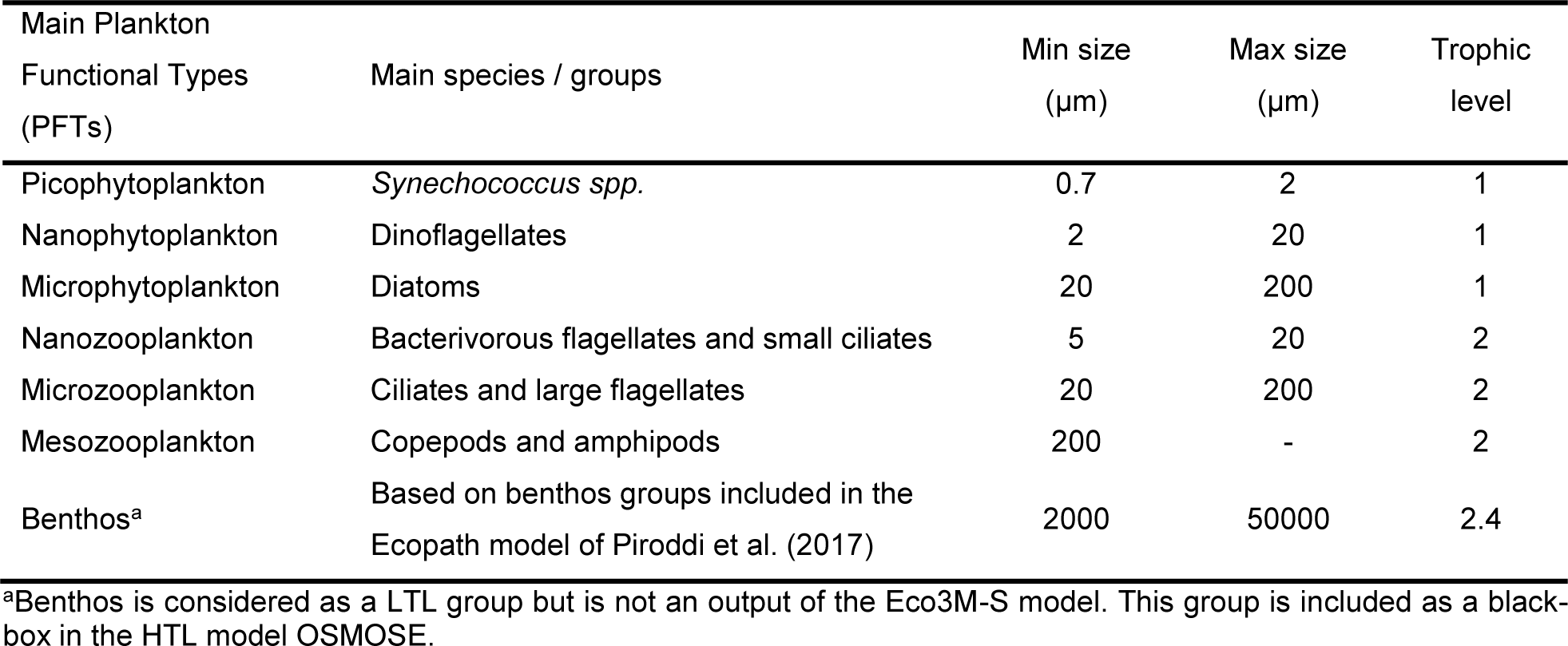
Parameters of the seven low trophic level compartments used to build the trophic links with OSMOSE. Other parameters used to run ECO3M-S are documented in Auger et al. (2011), Ulses et al., (2016) and Kessouri et al., (2017).

### 2.2 The high trophic levels model OSMOSE

The OSMOSE model has been applied in different ecosystem types such as upwelling ecosystems (Southern Benguela and Humboldt), temperate ones (West Coast Canada, Jiaozhou Bay), Mediterranean ones (Gulf of Gabes, Gulf of Lions) or subtropical ones (West Florida shelf) in order to assess the impacts of both fishing and climate change scenarios on marine food web functioning and species resilience (Fu et al., 2013; Grüss et al., 2015; Halouani et al., 2016; Marzloff et al., 2009; Travers et al., 2009; Xing et al., 2017). OSMOSE is a size-based trophic model that focuses on high trophic levels, mainly fish species. This multispecies and individual-based model is spatially explicit and represents the whole life cycle of several interacting marine species. From eggs to adult fish, major processes of the life cycle, i.e. growth, predation, reproduction, natural and starvation mortalities as well as fishing mortality are modeled step by step. Under computational time and memory constraints, rather than being truly individual-based, OSMOSE is based on “super-individuals”, as proxies for fish schools, defined as group of individuals sharing the same age, length, diet and spatial position and interacting with other schools in a two-dimensional grid. Species interact through predation in a spatial and dynamic way (Shin and Cury, 2004). The model is forced by species-specific spatial distribution maps which can vary interannually, seasonally, or depending on ontogenetic stages. OSMOSE allows the emergence of complex trophic interactions from two basic assumptions on predation process: for a given individual (a school), prey consumption depends on the spatio-temporal co-occurrence of the predator and its prey (in the horizontal and vertical dimensions) and is conditioned by size compatibility between a predator and its prey. Thus, unlike other trophic models such as Ecopath with Ecosim (Christensen and Walters, 2004), species dynamics and trophic structures are not modelled from pre-established trophic interactions between species: each fish can potentially be a predator or a prey, regardless of its taxonomy, but depending on size compatibility between a predator and its prey (Shin et al., 2004; Shin and Cury, 2001). A maximum and a minimum predator/prey size ratio are thus defined to rule predator prey interactions (Travers et al., 2009). To integrate a vertical dimension in the food web, accessibility coefficients are defined in the form of a prey-predator accessibility matrix that reflects possible mismatches or overlap between species vertical distributions and/or potential refugia allowing a certain proportion of a fish school to remain inaccessible to predation. At each time step, a predation efficiency rate can be calculated for each fish school (i.e. the food biomass ingested within a time step over the maximum ingestion rate), from which growth, starvation and reproduction rates are determined. In OSMOSE, the functions defining growth and mortality are deterministic. The main source of stochasticity comes from the species movement within their habitat and the order at which schools interact (through predation). Model details and equations are provided in Appendix A and on https://documentation.osmose-model.org/.

### 2.3 Parameterization of OSMOSE-MED

OSMOSE-MED covers the whole Mediterranean basin, from the Strait of Gibraltar to the Levant basin and from the Northern Adriatic Sea to the Southern Ionian Sea (Figure 1). It extends from approximately 26.9°N to 46.3°N in latitude and from approximately 5.6°W to 36.1°E in longitude. The Marmara Sea and the Black Sea are not included in the model. The OSMOSE-MED is built on a regular grid divided into cells of 20×20 km (for a total 6229 cells). Grid resolution was a compromise between the fine scale ecology of the modelled species and computation time limitations. The time step was set according to the spatial resolution. Here we adopted a time resolution of 15 days within which species were assumed to have access to the first layer of surrounding cells when foraging for prey.

**Figure 1.**
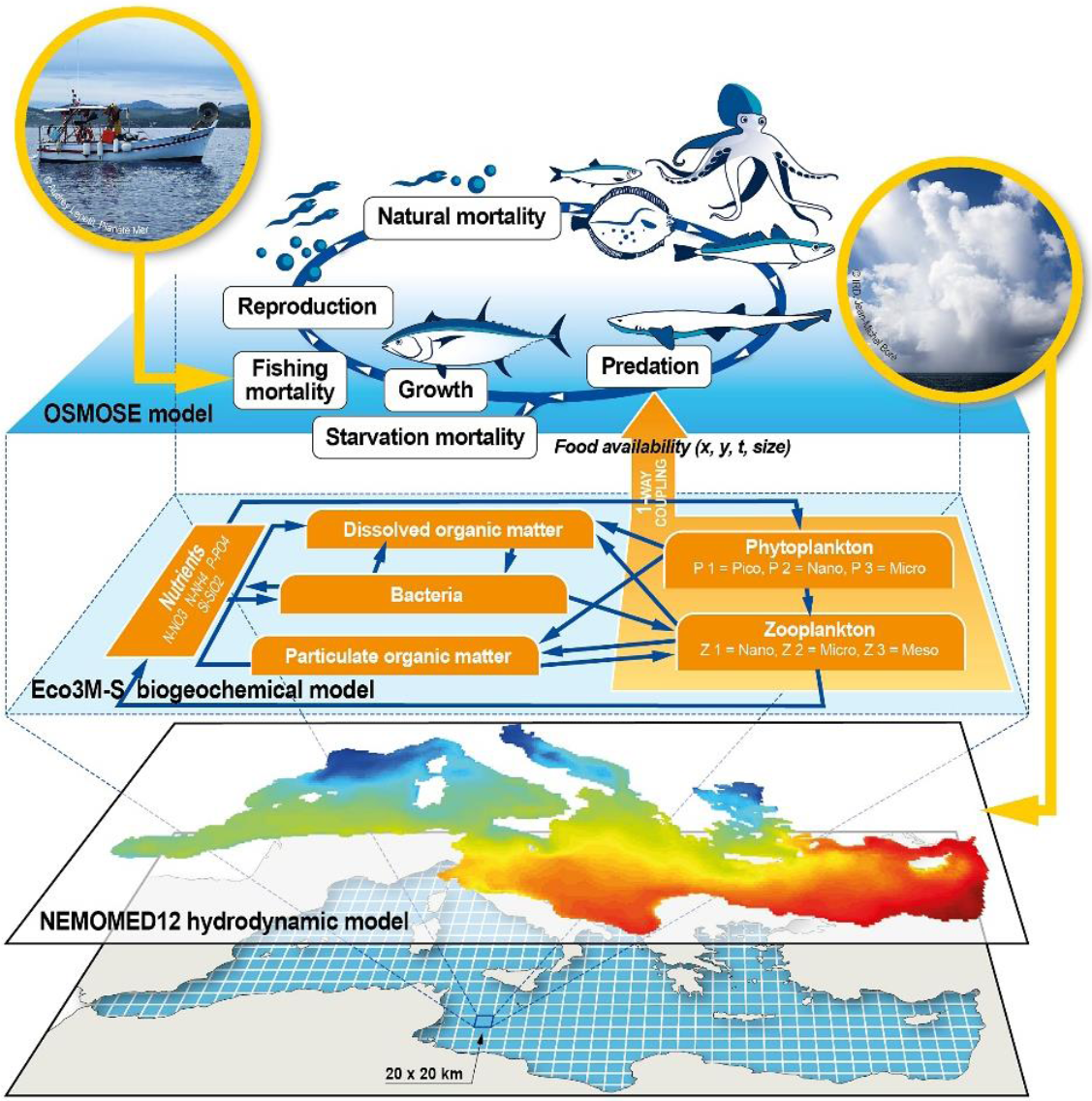
Conceptual representation of the End-to-End model “OSMOSE-MED” applied to the whole Mediterranean Sea. The high trophic levels OSMOSE model is forced (1-way coupling) by the biogeochemical Eco3M-S model through the predation by high trophic levels (i.e. fish, cephalopods and crustaceans) on low trophic levels (i.e. phyto and zoo-plankton). Eco3M-S is forced by the NEMOMED 12 hydrodynamic model. Impacts of climate variability and fishing mortality can be taken into account explicitly. (in color)

A 15-daily climatology was constructed from the 2006-2013 outputs of the biogeochemical model Eco3M-S and used to force the HTL model (offline coupling). The forcing model outputs were thus reflecting an average year in the period 2006-2013, characterized by a seasonal and a spatial variability of climate and plankton state variables. The coupling between NEMOMED 12/Eco3M-S and OSMOSE was realized through the predation process. At each time step and location, the biomass of the 6 plankton groups was used as potential prey fields forcing the HTL model. As within OSMOSE, predation upon plankton groups is an opportunistic size-based process (Travers-Trolet et al., 2014) controlled by a minimum and a maximum predation size ratio parameter. Benthic organisms (mainly invertebrates, crustaceans, polychaetes) are part of the diets of several HTL species included in OSMOSE-MED but they were explicitly modelled neither in ECO3M-S nor in OSMOSE. We therefore created an additional “benthos” compartment for which no life cycle or dynamics were modelled, but a few parameters were provided (size range and trophic level in Table 1) as well as a biomass level (derived from Piroddi et al. 2017) that was considered uniform over the Mediterranean Sea.

Regarding HTL species, 100 fish, cephalopod and crustacean species were explicitly modelled in OSMOSE-MED: 86 fish species, 5 cephalopods and 10 crustaceans (Appendix B). The selection of the 86 fish species was strongly dependent on data availability for model parameterization (biological parameters and life history traits for example) and for confronting model’s output to observations (species biomass data for example). Data search and mining for the parameterization of the modelled species life cycles represented a significant time investment. From the 635 fish species included in the FishMed database (Albouy et al., 2015), we searched the scientific literature and found the life history parameters (i.e. growth, reproduction and mortality) required to parameterize the OSMOSE model for only 86 fish species. Cephalopod and crustacean species were selected for their high commercial value, high contribution to total biomass and data availability. They also play an important role in food web dynamics (Peristeraki et al., 2005; Roberts, 2003) and represent key components in several Ecopath models applied to ecosystems in the Mediterranean Sea (e.g., Bănaru et al., 2013; Corrales et al., 2017; Hattab et al., 2013; Piroddi et al., 2017). All these species represent on average around 95 % of declared fisheries catches in the 2006-2013 period (FAO, 2006-2017). The biological parameters linked to growth (i.e. Von Bertalanffy parameters, length-weight relationship parameters), mortalities (e.g. maximum age, natural mortality not explicitly represented in OSMOSE, age/size at recruitment), reproduction (i.e. size at maturity, relative fecundity) and predation (minimum and maximum predation size ratios), along with their sources, are detailed in Appendix B and C. As much as possible, data were specific to Mediterranean ecosystems and were derived from or used by stock assessment working groups in the Mediterranean Sea.

Within each time step (15 days), the following events occur successively in OSMOSE-MED (Figure 1). First, each school is uniformly distributed over space according to a unique distribution map specified for each species (see 2.4). In this application of OSMOSE, due to the lack of observations, we did not account for any seasonal or ontogenic variation in fish distributions. As the maps do not change from one-time step to the next, schools can move to an adjacent cell or remain in the same cell following a random walk process (Shin et al., 2004; Travers-Trolet et al., 2014). Second, mortalities such as predation mortality, additional natural mortality and fishing mortality are applied to schools. The order at which schools interact as well as the order of mortality events is randomly drawn within each time step. Third, food intake, subsequent to predation events, modulates the growth in weight and size of species and their starvation level. Finally, reproduction occurs for fish having a length greater than the length at sexual maturity and allows the introduction of new schools of age 0 (eggs) in the system (Appendix A).

### 2.4 Modelling high trophic level species distribution

A niche modelling approach based on environmental data has been used to generate species distribution maps in the Mediterranean Sea. These distribution maps are then used in input of OSMOSE. Occurrences of the species included in OSMOSE-MED were compiled and merged from multiple sources: the Ocean Biogeographic Information System (OBIS: www.iobis.org), the Global Biodiversity Information Facility (GBIF: www.gbif.org) the Food and Agriculture Organisation’s Geonetwork portal (www.fao.org/geonetwork) and the atlas of Fishes of the Northern Atlantic and Mediterranean from the FishMed database (Albouy et al., 2015) (Appendix D). Values of environmental predictor variables at these locations were extracted from the World Ocean Atlas 2013 version 2 for climate data (https://www.nodc.noaa.gov/OC5/woa13/woa13data.html). To take into account the vertical distribution of species in the water column six environmental metrics were derived from monthly temperature and salinity climatologies: mean sea surface temperature and salinity (0-50 m depth), mean vertical temperature and salinity (0-200m depth) and mean sea bottom temperature and salinity (50m – maximum bathymetry depth). These metrics were used to model bioclimatic envelopes for each species. The use of environmental variables assumed that current species ranges are mainly driven by the abiotic environment, which is a reasonable hypothesis for marine species for which water temperature has been commonly considered as the main driver of fish geographic ranges (Ben Rais Lasram et al., 2010; Ben Rais Lasram and Mouillot, 2008; Cheung et al., 2009; Sabatés et al., 2006).

Present distributions were modelled using eight climate suitability models (generalized linear models, generalized additive models, classification tree analysis, boosted regression trees, random forests, multivariate adaptive regression splines, artificial neural networks and flexible discriminant analysis) embedded in the BIOMOD2 R package (Thuiller et al., 2009).

OBIS and GBIF databases provide only occurrence data at world scale (Hattab et al., 2014). To build reliable species distribution models, pseudo-absences (PAs) were generated in order to better characterize the environmental conditions experienced by species within their current ranges (Hattab et al., 2013b, 2014). PAs were selected randomly, outside the suitable area of the surface range envelope model. The number of simulated PAs was the double of occurrence data and they were equally weighted to the presence points during the fitting process.

In order to assess the accuracy of our final distributions, the True Skill Statistic (TSS, Allouche et al. (2006)) was used to measure the performance of each model. It represents a combined measure of the sensitivity (i.e. the proportion of correctly predicted presences) and specificity (i.e. the proportion of correctly predicted absences).

For each species, the consensus distribution was obtained with an ensemble forecast approach. Results were weighted according to the True Skill Statistic criterion (Allouche et al., 2006), i.e. weights were calculated on the basis of models’ accuracy in independent situations (Thuiller et al., 2009). To derive a consensus prediction, only the “best” model outputs (i.e. models with a TSS > 0.6) were kept (Appendix D). To transform the probabilistic consensus distribution into a presence/absence distribution, we preserved the occurrence probabilities for pixels above the sensitivity–specificity sum maximization threshold (i.e. the threshold that maximized the TSS criterion), and set to zero the occurrence probability for pixels under the threshold (Barbet-Massin et al., 2009). Spatial distribution maps are available in Appendix D.

### 2.5 Calibration of the end-to-end model OSMOSE-MED

An Evolutionary Algorithm (EA), inspired by the process of Darwinian evolution and developed for the calibration of complex stochastic models such as OSMOSE, has been used to calibrate the E2E model OSMOSE-MED (Duboz et al., 2010; Oliveros-Ramos and Shin, 2016). By estimating some unknown parameters (i.e. larval mortality rates of HTL species, availability coefficients of LTL species to all HTL species and fishing mortality for exploited species), the calibration process aims to constrain predicted biomasses and catches of HTL species by OSMOSE-MED within realistic ranges. The model was confronted to observed data using a maximum likelihood approach (Oliveros-Ramos et al. 2017). A log-normal distribution was assumed for the biomass and catch errors.

The aim of the EA is to optimize an objective function over a given search parameter space, a penalized negative log-likelihood function in our case (Oliveros-Ramos et al., 2017). A population of ‘individuals’, where each individual is a set of parameters (called the genotype) in the search space, is first created. Different unknown combinations of parameters are tested in order to minimize the objective function. Computation of the phenotype (i.e. outputs produced by a run of OSMOSE-MED with a given set of parameters) and of the fitness (i.e. goodness of fit from the minimization of the negative log-likelihood function) is done in a second step. At each generation (i.e. iteration of the optimization process), the algorithm calculates an “optimal parent”, which results from the recombination of the parameter sets that provide the best solution for each objective (partial likelihoods for species biomass and catch) (Oliveros-Ramos and Shin, 2016). The optimal parent is then used to produce a new set of parameter combinations (by recombination/mutation) which constitutes the next generation. The EA is run until the convergence of the objective function or is stopped after a given number of generations (Duboz et al., 2010; Oliveros-Ramos et al., 2017; Oliveros-Ramos and Shin, 2016).

A steady state calibration of the OSMOSE-MED model was performed using the mean of reported and reconstructed catches averaged over the period 2006-2013 (called hereafter “reference state period”) as target data. For tuna and other large pelagic species (e.g., the sworfish *Xiphias gladius*), catch data were extracted from ICCAT statistics database. For all other exploited species, reported fisheries landings were provided by the FAO-GFCM database (http://www.fao.org/gfcm/data/capture-production-statistics) and reconstructed catches were obtained from the Sea Around Us project (Zeller and Pauly, 2015). Reconstructed catches from the Sea Around Us were used in order to reduce data gaps and take into account discards and illegal, unreported and unregulated fishing in the Mediterranean Sea, where actual catches are often underestimated (European Commission, 2003; Moutopoulos and Koutsikopoulos, 2014).

Cumulated biomass from stock assessments from different GSAs (Geographical Sub-Area) of the Mediterranean Sea were used when available and realistic (i.e. when cumulated available biomass by species was higher than the average of FAO-SAU catches), and averaged over the reference state period (e.g., for *Merluccius merluccius, Sardina pilchardus or Engraulis encrasicolus*) (Appendix C). Biomass estimates of *Thunnus thynnus* and *Thunnus alalunga* were based on experts’ knowledge (Fromentin J.M. and Winker H., personal communications). For all other species for which biomass estimates were not available, we applied strong penalties to the objective function when output biomass from OSMOSE-Med did not stand within plausible ranges. Specifically, we considered FAO reported catches as a minimum threshold for species biomass and the maximum biomass threshold was derived from mean FAO-SAU catches and a fisheries exploitation rate of 15 % that is assumed to be a very low exploitation rate in the context of Mediterranean fisheries (Vasilakopoulos et al., 2014),

The model was run for 100 years for each set of parameters to make sure that OSMOSE-MED reached a steady state and only the last 30 years were analyzed by the EA. The calibration process allowed to estimate a set of parameters for each species represented in the OSMOSE-MED: coefficients of plankton accessibility of the 7 LTL groups considered in the model (7 parameters), larval mortality rates of the 100 HTL species (100 parameters) and fishing mortality rates for species for which catch data were available (87 parameters).

Following the methodology described in Oliveros-Ramos et al. (2017), a sequential multi-phase calibration was applied to estimate the 194 unknown parameters (Oliveros Ramos, 2014), with three successive calibration phases detailed in Table 2.

**Table 2.**
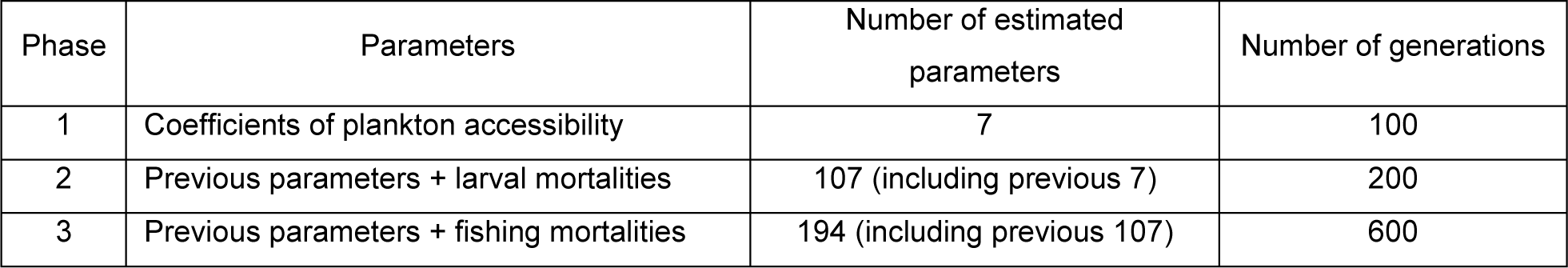
Order at which parameters were estimated in the multi-phase calibration of the OSMOSE-MED model, using the evolutionary optimization algorithm included in the Calibrar R package.

The optimization process involved the use of Calibrar and OSMOSE R packages (Oliveros-Ramos et al., 2017; Oliveros-Ramos and Shin, 2016) available from the CRAN website (https://cran.r-project.org/web/packages/calibrar). The calculation was performed using DATARMOR, the High Performance Computing (HPC) facilities of “Pôle de Calcul et de Données Marines” of IFREMER (https://wwz.ifremer.fr/pcdm/Equipement). 36 compute nodes representing 1008 cores (2.4Ghz) and around 4 To of RAM (Random Access Memory) were mobilized to perform the calibration, which involved several iterative trials during more than one year.

Due to the inherent stochasticity of OSMOSE, 10 replicated simulations (i.e. with identical set of parameters) were averaged to analyze the outputs of the last 10 years.

### 2.6 Evaluation of OSMOSE-MED outputs with independent data

In order to evaluate the capacity of OSMOSE-MED to predict the spatial distribution of the whole biomass in a realistic way, we confronted the model’s output to observed data that were not used neither for the calibration of OSMOSE-MED, its parameterization, nor the climate niche modelling that led to species distribution maps. The ranking of Geographical Sub-Areas (GSAs), based on cumulated biomass estimates by species (in kg.km^-2^) from the MEDITS survey (International bottom trawl survey in the Mediterranean, Bertrand et al., 2002) in 2006-2013 was compared to the ranking predicted from OSMOSE-MED (see Appendix F for the correspondence between GSA number, their names and their sizes). To evaluate the consistency of the OSMOSE-MED model at the community level, the mean trophic level (mTL) of each species has been calculated and compared to three different sources: the FishMed database, a database which contains ecological and biological traits for 635 Mediterranean fish species (Albouy et al., 2015), the Ecopath model built at the Mediterranean basin scale by Piroddi et al. (2017, 2015a) and a review of feeding habits and trophic levels of 148 Mediterranean fish species (Karachle and Stergiou, 2017; Stergiou and Karpouzi, 2002).

An important step in the validation of the model lied in the confrontation of simulated species diets to observations and to the current knowledge of the trophic functioning of the Mediterranean ecosystem. In OSMOSE, the diet composition of a species is not determined *a priori* in input of the model but it emerges from the assumption of an opportunistic predation process, based on predator-prey size constraints and spatio-temporal co-occurrence. To check whether this size-based predation rule led to realistic and consistent dietary features, we focused on the diet compositions of four of the most important species in terms of volume or value of catches in the Mediterranean Sea, namely the European anchovy, the European pilchard, the Red mullet and the European hake. We specifically compared the diets of the adults from OSMOSE-MED to the diets derived from the mass-balanced Ecopath model of the Mediterranean Sea (Piroddi et al., 2015a) as the functional groups in the latter model were mostly parameterized to represent the adults. The diet matrix used for parameterizing Ecopath was compiled from the available literature and mostly based on empirical data (Piroddi et al., 2015a, 2017) it was thus used as a convenient way to access observed diets and current knowledge on major trophic interactions, at least for the well-studied species.

## 3 Results and discussion

### 3.1 Calibration

OSMOSE-MED reached a steady state after around 50 years of simulation. The evolutionary algorithm converged and stabilized after 500 generations. Both negative log-likelihoods and global AIC improved during each phase but regarding the global evolution of the likelihoods larval mortalities seemed to be the parameters playing the most important role in the calibration process. Accessibility coefficients of LTL groups to HTL organisms ranged between around 10^-9^ and 10^-1^ (Appendix E). The smallest values were obtained for small size plankton groups (except for picophytoplankton), which could be expected in view of their high biomass and low predation rates by HTL organisms (Jackson and Lenz, 2016; Morote et al., 2010; Pepin and Penney, 2000). In contrast, higher coefficients were found for mesozooplankton and benthos groups, for which around 1 and 0.5 % were respectively available to predation by HTL. These coefficients were in the same order of magnitude than in other modelled ecosystems (e.g Grüss et al., 2015; Marzloff et al., 2009; Travers-Trolet et al., 2014).

Estimated larval mortality rates (*M*_*0*_) ranged between 0.14 and 10.60 year^-1^ for the caramote prawn (*Penaeus kerathurus*) and the small-spotted catshark (*Scyliorhinus canicula*), respectively (Appendix E). The larval mortality rate found for *P. kerathurus* is probably too low compared to the value (*M*_*0*_ *=*1.58 *year*^*-1*^) estimated by Halouani et al. (2016b) with the OSMOSE-GoG model and the biomass estimated by our model stands outside a valid interval. The majority of larval mortalities were comprised between 1.49 and 5.29 *year*^*-1*^ (mean = 3.69 ± 2.70 *year*^*-1*^; Appendix E). A low larval mortality rate estimated by the evolutionary algorithm for a particular species does not necessarily mean that the total natural mortality is small but may also reflect that most of the sources of mortality (predation by the other modelled species for example) are simulated explicitly in the model (Travers-Trolet et al., 2014).

Because fishing mortality rates (*F*) estimated by stock assessments were not available for all exploited species, we have chosen to estimate these parameters by confronting the model’s output to observed and reconstructed catches during the third phase of the calibration process. Most of the fishing mortality rates were within the range of 0.23 to 0.8 *year*^*-1*^ and global fishing mortality rate was on average 0.60 ± 0.48 *year*^*-1*^ (Appendix E).

### 3.2 Confronting OSMOSE-MED to observations and current knowledge

#### 3.2.1 Species biomass

The estimated biomass, averaged over the last ten years of a simulation and over ten replicates, were globally in acceptable intervals (i.e. above FAO reported catch and below a theoretical maximum biomass if we consider an exploitation rate of 15 % for the averaged FAO-SAU catches) (Figure 2). For species for which biomass from stock assessments were available, as is the case for instance for the European pilchard (*Sardina pilchardus*), the European anchovy (*Engraulis encrasicolus*) or the European hake (*Merluccius merluccius*), the total biomass predicted by OSMOSE-MED were slightly higher or very close to previously estimated biomass (Figure 2). Given that stock assessments were mostly available for European waters, a higher estimated biomass for species like *Sardina pilchardus, Parapenaeus longirostris* or *Mullus barbatus barbatus* could actually reflect a biomass volume present in the southern part or in areas not assessed in the Mediterranean Sea. Overall, the European anchovy and the European pilchard (around 1.8 millions of tons of biomass) represented, in cumulated, around 50 % of the total biomass of the system (excluding plankton organisms). The prevalence, in terms of biomass, of pelagic fishes was also found in an Ecopath model of the Mediterranean Sea (Piroddi et al., 2015a). For species like *Crangon crangon, Atherina boyeri* and *Etrumeus teres*, due to their highly variable population dynamics (high fecundity, short lifespan, high biomass turnover rate), biomass have been particularly difficult to calibrate and were finally overestimated by OSMOSE-MED. Moreover, for non-native species (e.g. *Etrumeus teres*), more research is needed on their biology and ecology in their new expansion areas, in order to get robust life history traits estimates, and to improve model predictions (Dimarchopoulou et al., 2017; Katsanevakis et al., 2014, 2012). The lack of stock assessments and the sometimes difficult access to the results of these assessments constitute some real barriers to the development, parameterization and calibration of ecosystem models in the region (Coll et al., 2013; Katsanevakis et al., 2015; Piroddi et al., 2015a). Around 25 % of landed biomass and less than 10 % of exploited stocks are currently assessed and on an irregular basis (Tsikliras et al., 2015). Moreover, the monitoring of fish stocks is hindered by the lack of biological or ecological observational data for far too many stocks, with approximately 80 % of landings coming from data-deficient stocks (Dimarchopoulou et al., 2017; Le Quesne et al., 2013).

**Figure 2.**
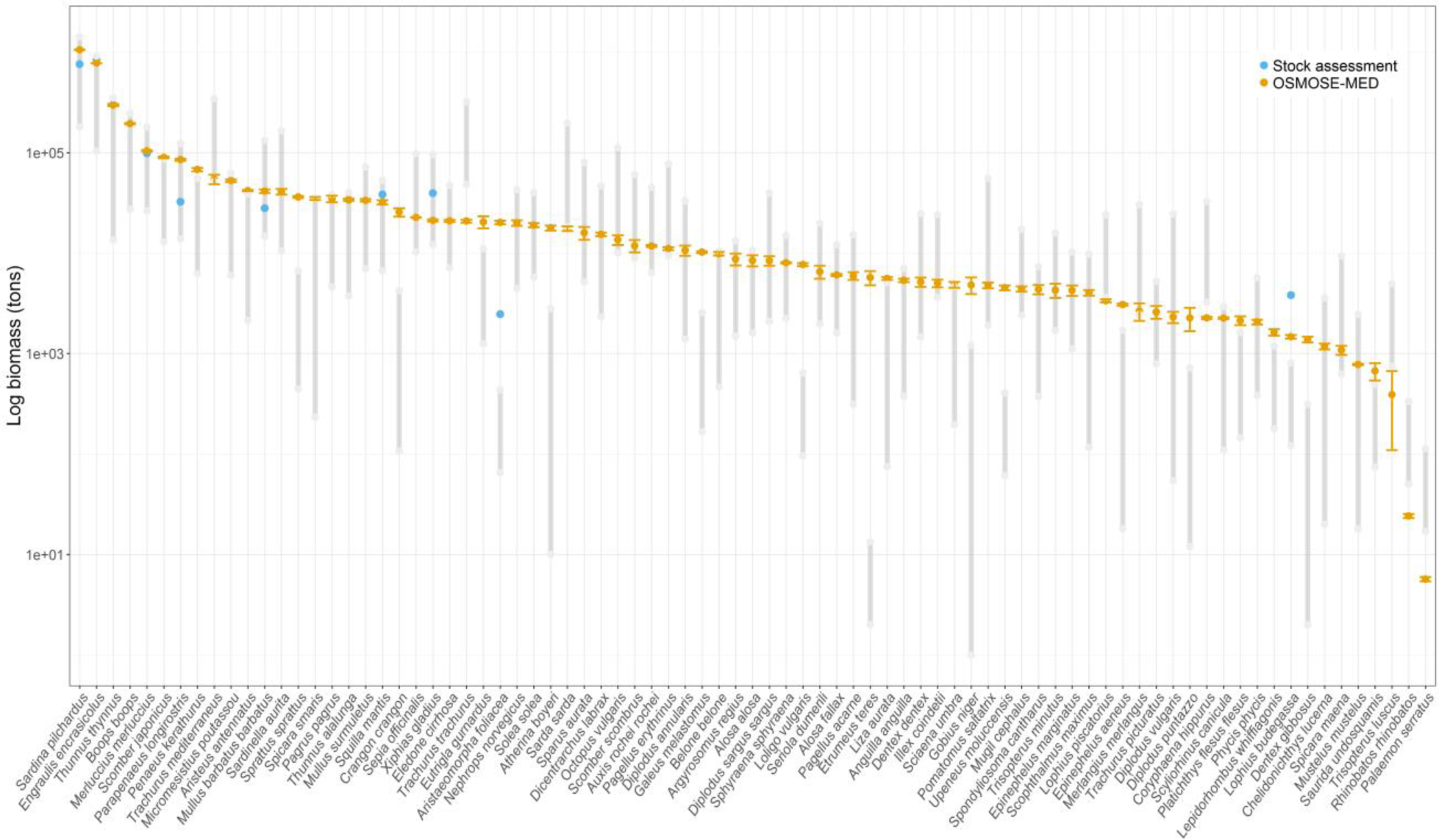
Average predicted biomass in log scale and associated standard deviation of exploited species (87 species out of the 100 modelled) (orange circles). Blue circles represent cumulated biomass from stock assessments (only cumulated biomass higher than FAO reported catch were represented). Minimum and maximum of grey segments are respectively the FAO reported catch and a theoretical maximum biomass considering an exploitation rate of 15 % and the average between FAO and Sea Around Us catch. (in color)

The model properly predicted the spatial distribution of the overall biomass, at least for the northern part of the Mediterranean Sea where MEDITS survey were conducted, as suggested by the significant Spearman’s rank correlation coefficient value of 0.71 between MEDITS and OSMOSE-MED biomass ranking. Existing differences between the ranking of some GSAs can be explained in two ways. For instance, in Corsica Island, OSMOSE-MED predicted less relative biomass (rank 15 out of a total of 16 GSAs) than estimated by MEDITS survey (rank 8). This is partly due to the very narrow continental shelf around Corsica Island and to the resolution of our model (20×20 km^2^) that could be too coarse to represent the dynamics in this area. Therefore, the climate niche models and resulting distribution maps in input of OSMOSE-MED were not resolving precisely enough the spatial distribution of the species closely associated to the Corsican continental shelf. The development of OSMOSE-MED at a finer resolution scale has been attempted in the early stages of the model configuration, but the computational cost has been judged too high for the calibration process (at least two to three times the necessary computation time for a 10×10km^2^ resolution). On the contrary, for GSAs ranking higher in OSMOSE-MED than in MEDITS ranks (i.e. below the 1:1 line in Figure 3), the differences could be explained by the fact that MEDITS is a demersal trawl survey thereby having a low catchability for small pelagic fishes. Even though trawl survey data were useful for assessing the spatial and temporal trends of pelagic species in the Mediterranean Sea (Brind’Amour et al., 2016), some biases may exist such as the biomass of some small pelagic fishes being potentially largely underestimated by the survey.

**Figure 3.**
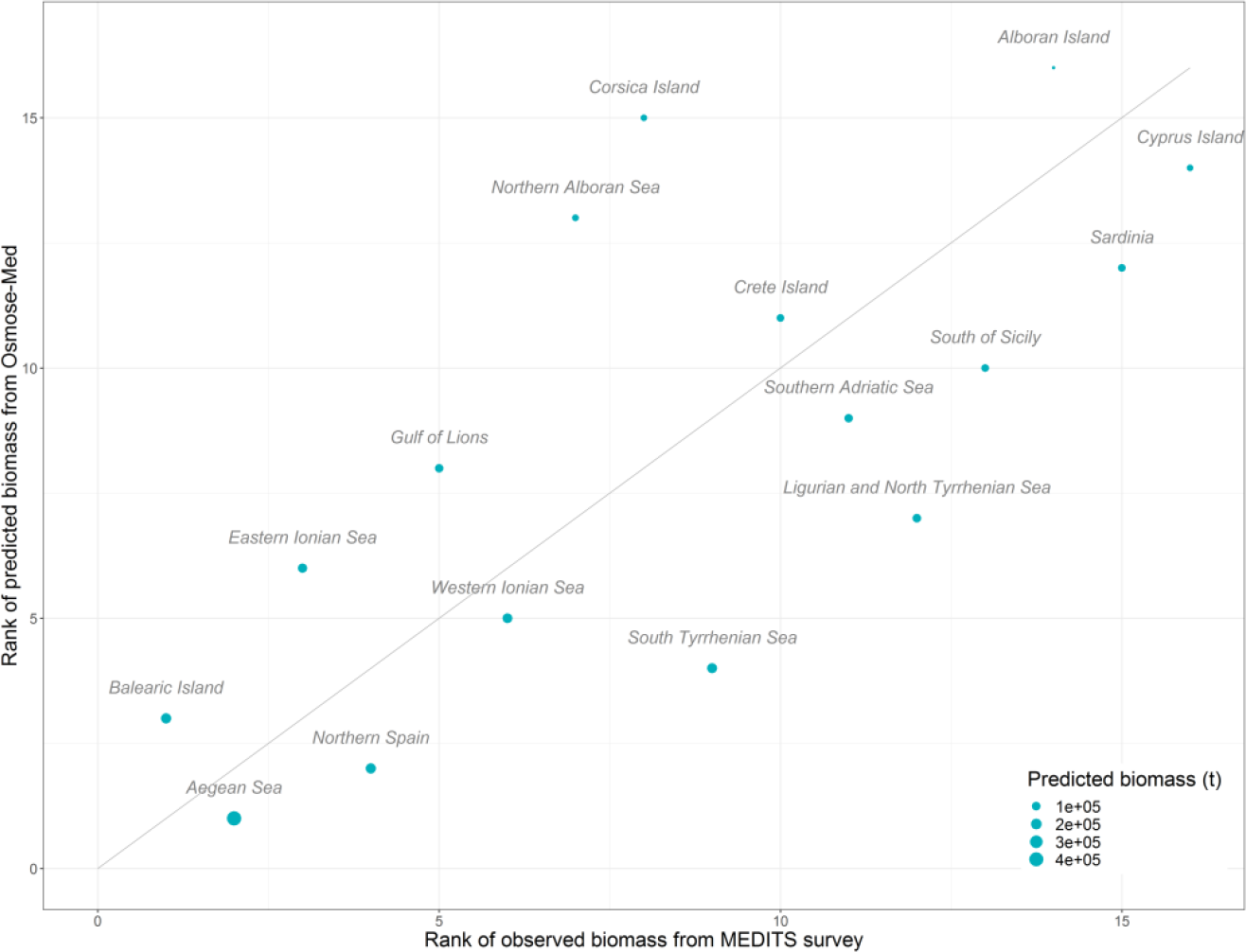
Comparison of predicted and observed ranks of total biomass by Geographical Sub-area (GSA). Observed total biomass are from the MEDITS survey (2006-2013). Circle size is proportional to the total predicted biomass by GSA. Solid line is 1:1 relationship. (in color)

#### 3.2.2 Species catches

Catches predicted by OSMOSE-MED were globally consistent with catch data in the Mediterranean Sea (Figure 4 and Figure 5). Our model predicted a total catch of around 802 470 t at the whole basin scale that compares well with the 681 243 t recorded by the FAO and with the 952 930 t reconstructed by the Sea Around Us (817 087 t in average). The European pilchard and the European anchovy represented almost 30 % of the total catches in OSMOSE-MED and around 40 % in reported or reconstructed catches over the 2006-2013 period (FAO, 2016; Pauly and Zeller, 2016). According to Stergiou et al. (2015), small pelagic species, mainly European anchovy and European pilchard, dominate the landings across the entire Mediterranean Sea, with 34 % of cumulated landings in the western Mediterranean, 41 % in the central part and 25 % in the eastern part. The Spearman’s correlation coefficient between the rank of the average FAO-SAU catches by species and that estimated by OSMOSE-MED was 0.79 (Figure 5). The main differences between predicted and averaged reported-reconstructed catches came from the under-or over-estimation of species biomass by the model. For instance, the common prawn (*Palaemon serratus*) seemed to be underestimated in terms of predicted biomass and catch. For species for which biomass estimated by stock assessment were available, the OSMOSE-MED model predicted the catches relatively well. The estimated catches for the European anchovy was, for instance, around 118 480 t in OSMOSE-MED while reported and reconstructed catches by the FAO and the SAU over the 2006-2013 period were 103 650 t and 169 870 t, respectively.

**Figure 4.**
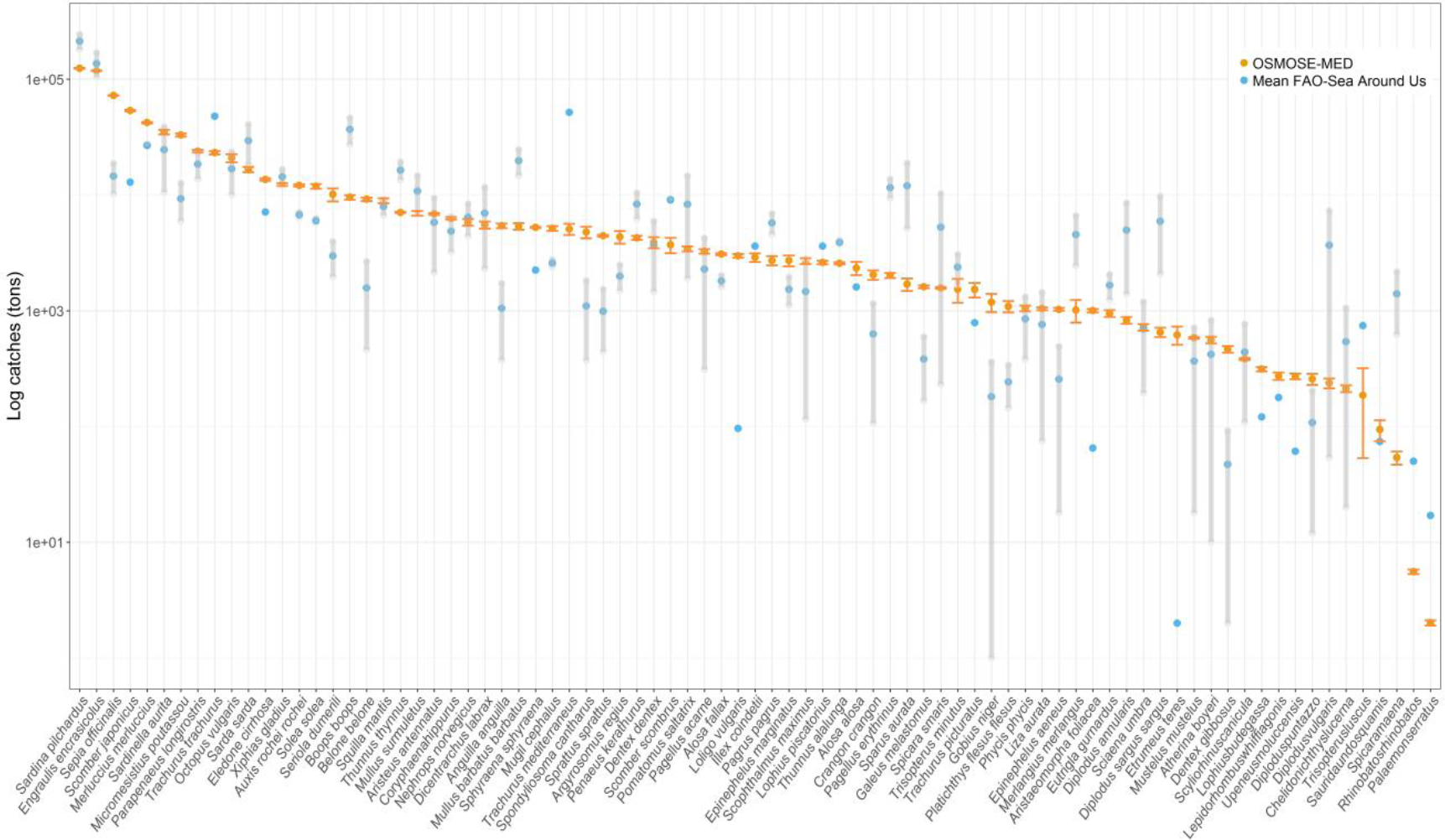
OSMOSE-MED average predicted catches on log scale and associated standard deviation of all exploited species (orange circles). Blue dots represent the average FAO – SAU catch which served as target data during the calibration process. Minimum and maximum of grey segments are the FAO reported catch and the SAU reconstructed catch, respectively. Predictions and data for the 2006-2013 period. (in color)

**Figure 5.**
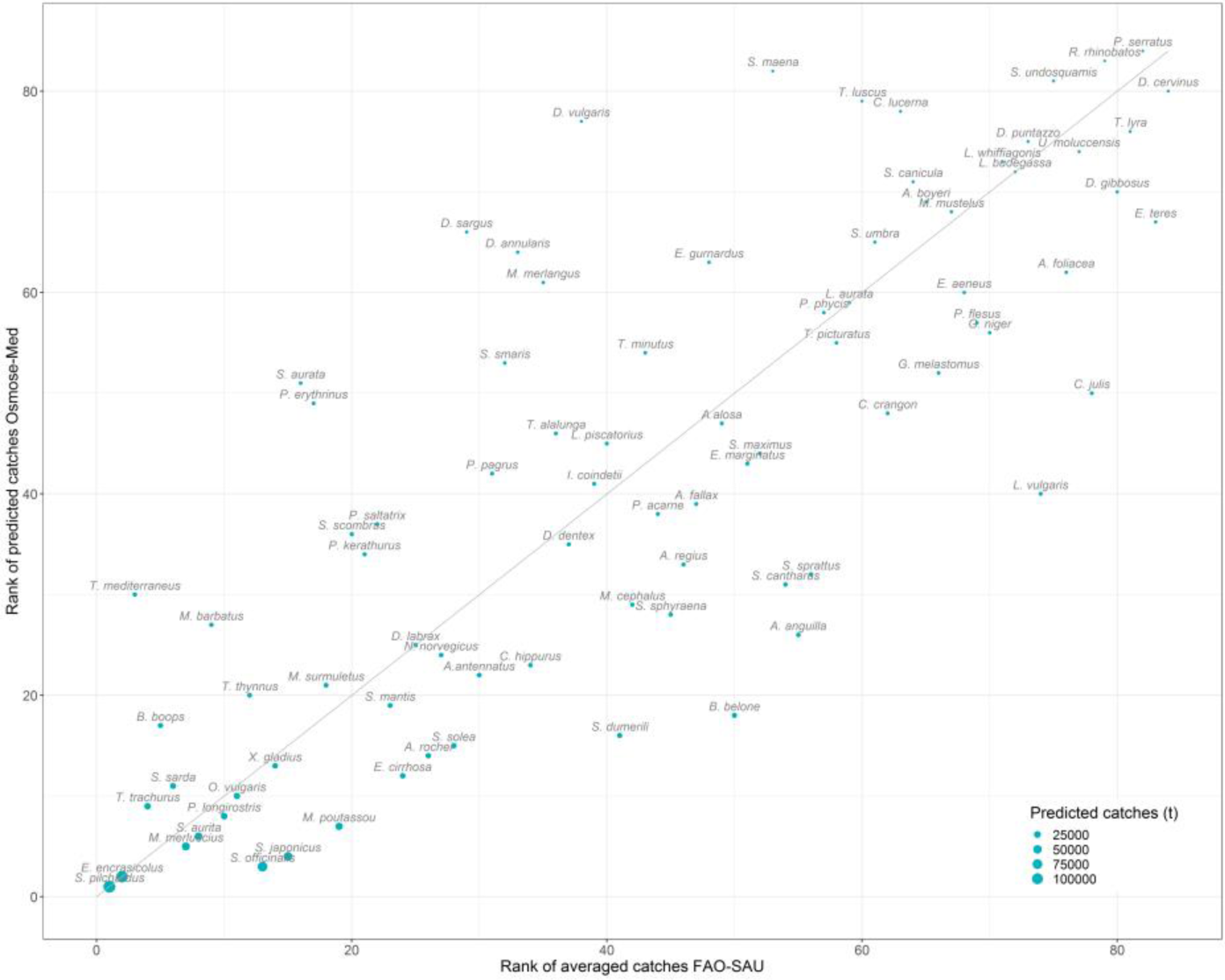
Comparison of predicted and observed (average FAO-SAU catches) ranks of catches by species. Circle size is proportional to the predicted catches. Predictions and data for the 2006-2013 period. Solid line is 1:1 relationship. (in color)

In the present version of OSMOSE (Version 3 update 2), fishing effort is homogeneous in space. Catch outputs could be improved with a spatialization of the fishing effort which is implemented in the last version under development. However, in the Mediterranean Sea, data on fishing effort and distribution is either unavailable or difficult to access in some regions (Katsanevakis et al., 2015). One solution could be the use of the new “Global Fishing Watch” database which collects data from the automatic identification system (AIS) of fishing fleets all around the world (Kroodsma et al., 2018). An index of the fishing effort in the Mediterranean Sea could be calculated by evaluating the fishing time by vessel characteristics (Kroodsma et al., 2018). However, as most of the vessels composing the Mediterranean fleet are less than 10 m and AIS is only compulsory for large European vessels, estimations would remain underestimated (Ferrà et al., 2018). Fitting an ecosystem model based on catch data is a difficult task in the Mediterranean Sea due to the poor quality of fisheries statistics (Pauly et al., 2014; Piroddi et al., 2017). A significant quantity of catches is still not recorded or some stocks are data-deficient. The almost twice difference between reported and reconstructed catches highlighted by Pauly and Zeller (2016) illustrates this issue. As suggested by Piroddi et al. (2017), better catch data and improved availability for modelling studies could help to estimate more realistic fishing mortalities and trends in space and time. The new MedFish4Ever initiative, launched by the European Commission in 2017 to rebuild a sustainable fisheries sector, could play a key role in the improvement of such data, at least for the northern Mediterranean (https://ec.europa.eu/fisheries/inseparable/en/medfish4ever).

#### 3.2.3 Species trophic levels

In general, the trophic levels from OSMOSE-MED were consistent with the results obtained by other studies in the Mediterranean Sea (Figure 6). 69 % of the OSMOSE-MED mTLs were close to previously estimated mTLs by less than 0.3. Among the 81 species that had several mTL data sources, OSMOSE-MED mTLs stood within the range of previously estimated mTLs for 58 species (72 % of the species). Trophic levels from OSMOSE-MED were generally higher than those of the Ecopath model and generally lower than those of FishMed which were mainly taken from the Fishbase database (Albouy et al., 2015). The significant Spearman’s correlation coefficients between the trophic levels from OSMOSE-MED and the trophic levels coming from FishMed, Ecopath and the review of Karachle and Stergiou (2017) were 0.67, 0.51 and 0.68, respectively. In OSMOSE-MED, Swordfish *Xiphias gladius* had the highest trophic level (mTL = 4.64 ± 0.002) whereas European pilchard had the lowest (mTL = 3.11 ± 0.0003). Large pelagic fish species like Swordfish, Dolphinfish (*Coryphaena hippurus*), Bluefin and Albacore tunas (*Thunnus thynnus and Thunnus alalunga)*, Atlantic bonito (*Sarda sarda*), shark species like Common guitarfish (*Rhinobatos rhinobatos*), Common smooth-hound (*Mustelus mustelus*) and Small-spotted catshark (*Scyliorhinus canicula*) and demersal species such as European hake (*Merluccius merluccius*) were all identified as top-predators in the OSMOSE-MED model (i.e. mTL > 4.15). These results are consistent with other trophic models built in the Mediterranean Sea which identified large pelagic fish species and sharks species (except from Common guitarfish) at the top of the food web (Albouy et al., 2010; Coll et al., 2007; Corrales et al., 2015; Halouani et al., 2016; Hattab et al., 2013a).

**Figure 6.**
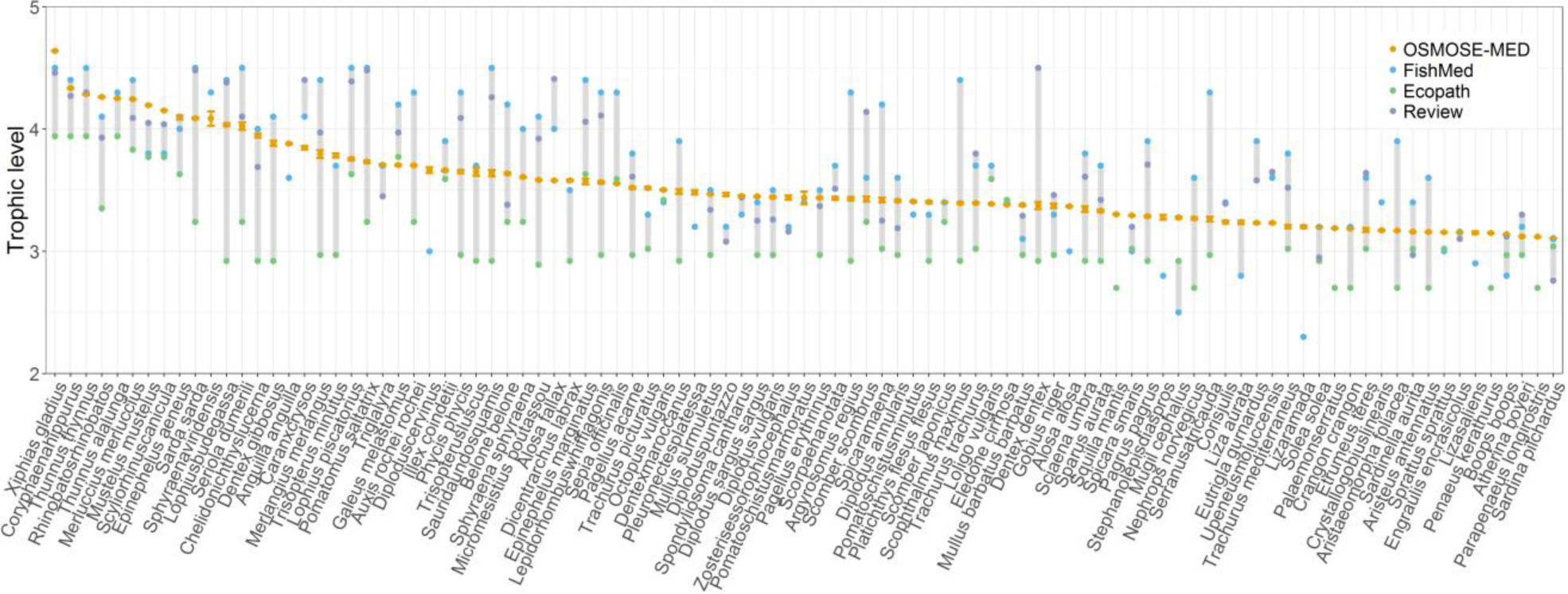
Mean predicted trophic levels of OSMOSE-MED species (orange circles) and trophic levels from the FishMed database of Albouy et al. (2015) (blue circles), the Mediterranean Ecopath model from Piroddi et al. (2017, 2015a) (green circles) and a review proposed by Karachle and Stergiou (2017) (purple circles). (in color)

#### 3.2.4 Species diets

OSMOSE-MED and the Mediterranean Ecopath model were more or less in agreement with regard to the prey composition of the diet of the four species under scrutiny (Figure 7). For European anchovy and the European pilchard, the simulated diets were similar and largely dominated in composition by the zooplankton, a pattern which is in agreement with other observations (Karachle and Stergiou, 2017; Stergiou and Karpouzi, 2002). In OSMOSE-MED, European pilchard consumed less phytoplankton (4.5 %, mainly diatoms) than in the Ecopath model (10 %) but the result remains qualitatively realistic (i.e. the main prey is zooplankton followed by phytoplankton). The dominance of zooplankton prey in the diet of pilchards could be explained in two different ways. Firstly, the availability coefficients of phytoplankton groups to HTL organisms were estimated to be very low by the model calibration (range between 10^-1^ to 10^-7^), which does not allow European pilchard to feed more on these groups. Secondly, it has been shown that populations of European pilchard, living in lower productivity region, as is the case for the Mediterranean Sea, would preferentially capture larger individual prey via particulate feeding and would consume more zooplankton than populations of the Northwest Atlantic (Costalago et al., 2015). Regarding Red mullet (*Mullus barbatus barbatus*) the main difference between the two models lies in the higher proportion of zooplankton preyed in OSMOSE-MED. Some of the crustaceans eaten in the Ecopath model were either included in the benthos group in OSMOSE-MED or explicitly modelled at the species level as is the case for *P. longirostris* and *P. kerathurus* that appeared in the simulated diet of Red mullet. For European hake, most of the prey eaten in OSMOSE-MED were grouped in more aggregated trophic boxes in Ecopath. For instance, shrimps were grouped in the functional group “crustaceans” in Ecopath, Octopus were grouped in the “benthic cephalopods” compartment and some species like *Mullus surmuletus* or *Boops boops* were grouped in the “small demersals” functional group. However, the percent contribution of some prey like European pilchard or European anchovy differs between the two models. The European pilchard represented, for instance, 5.7 % of the diet of the European hake in OSMOSE-MED and 12.5 % in Ecopath. However, hake diet varies greatly as a function of prey availability and abundance, in Mediterranean Sea as well as in other areas of the Atlantic Ocean (Carrozzi et al., 2018; Cartes et al., 2009; Velasco and Olaso, 1998). Carrozzi et al. (2018) found, for instance, that in the central Mediterranean Sea, European pilchard and European anchovy represented 3.78 and 1.32 % of hake diet, respectively.

**Figure 7.**
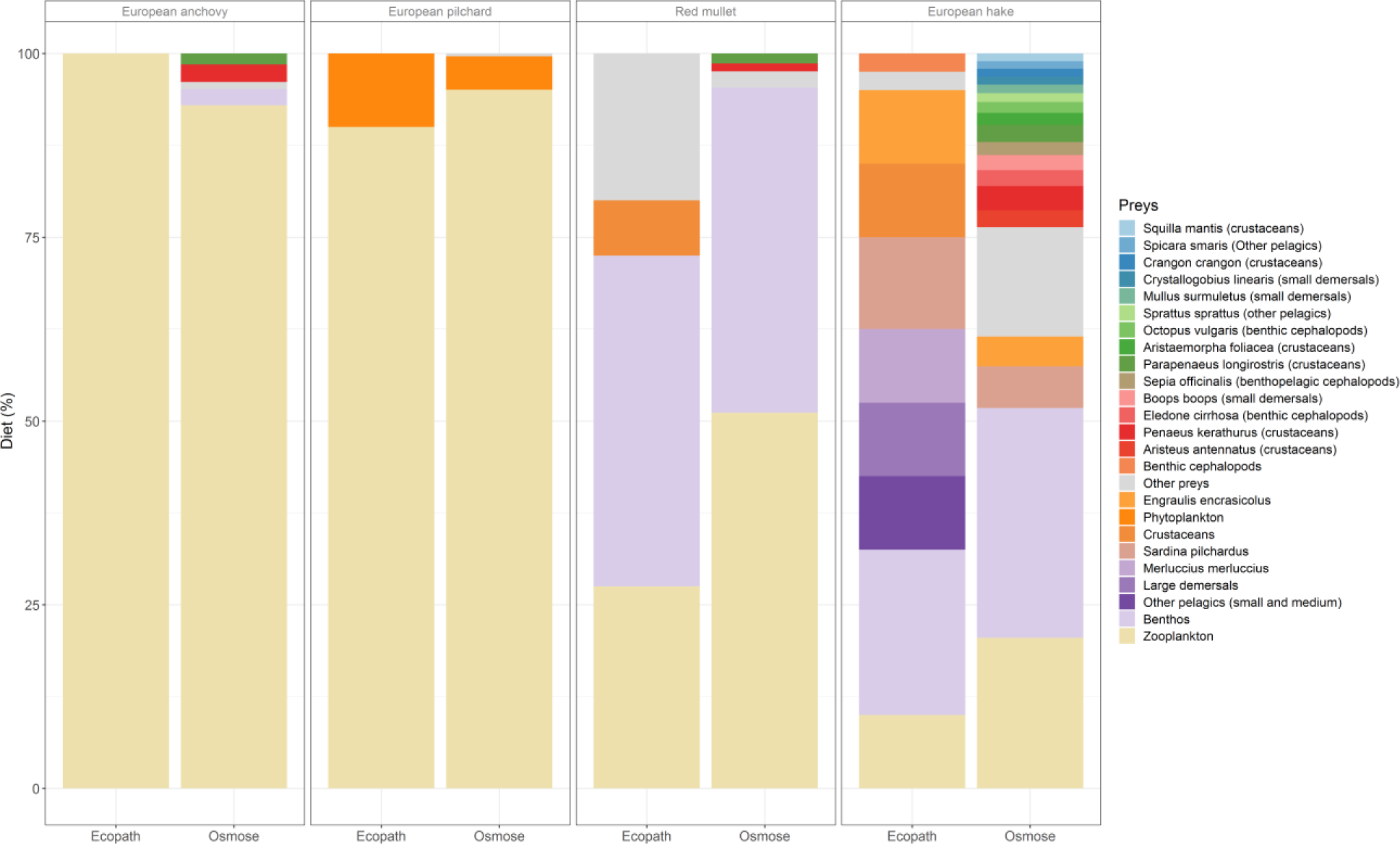
Diets simulated by OSMOSE-MED and the Mediterranean Ecopath model for four species (two small pelagic fish species (European anchovy and European pilchard) and two dermersal fish species (Red mullet and European hake)). Functional groups of Ecopath model in which OSMOSE-MED species are grouped are indicate in brackets. In both cases diets are expressed as percentage of prey by mass. (in color)

#### 3.2.5 Emerging spatial patterns

The total biomass (all HTL species confounded) was mainly distributed on the continental shelf and in areas where the primary and secondary productions were higher (Figure 8) in line with past studies (Durrieu de Madron et al., 2011, Bosc et al. 2004). The higher biomass found in highly productive areas (the Gulf of Lions, the Catalan Sea or the South Levantine Sea with the Rhône, the Ebro and the Nile rivers enhancing primary productivity through nutrient discharge, hence playing a major role for local food webs) suggested that primary production, by bottom-up control, was one of the main drivers of the biomass distribution of HTL organisms in the Mediterranean Sea. Numerous Ecopath models built at more local scales in the region confirm this assumption (Coll et al., 2006, 2007; Coll and Libralato, 2012; Halouani et al., 2016; Hattab et al., 2013a). The control of marine productivity, from plankton to fish, principally mediated through bottom-up processes that could be traced back to the characteristics of riverine discharges has also been demonstrated by Macias et al. (2014). These characteristics render the Mediterranean Sea vulnerable to sources of potential impacts on primary production such as climate change or marine pollution (Cheung et al., 2011; Jochum et al., 2012; Macias et al., 2015; Moullec et al., 2016) and highlight the need of expliciting the forcing of physicochemical oceanographic drivers on the dynamics of high trophic level organisms in a single modelling framework, to deal with possible bottom-up control and improve our capacity to predict future ecosystem changes (Piroddi et al., 2017; Rose et al., 2010; Travers-Trolet et al., 2014). On the other hand, since fishing effort was spatially uniform in our model, we could not assess precisely the direct role of fishing in the spatial distribution of the HTL biomass but rather the impacts on species biomass, species composition and interactions which were indirectly reflected by the biomass distribution over the Mediterranean Sea.

**Figure 8.**
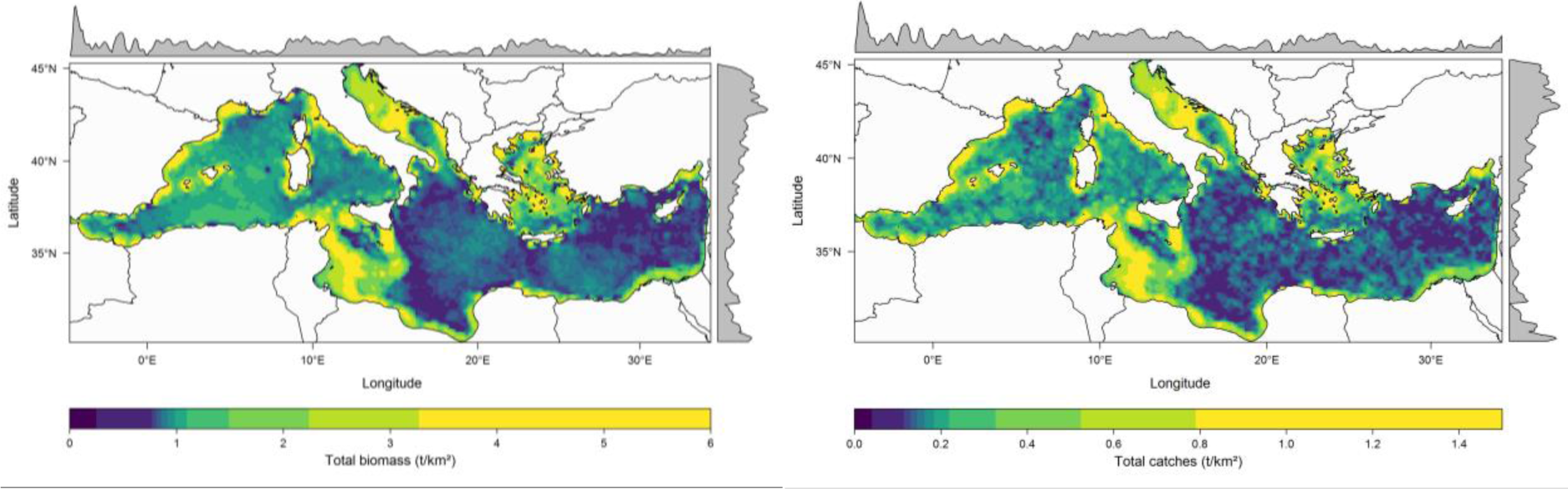
Spatial distribution of the simulated total biomass (left) and catches (right) (all HTL species confounded) expressed in t.km^-2^. Upper and right hand side plots respectively represent the meridionally and zonally averaged distribution of biomass and catches. (in color)

A low gradient of biomass is observed from Northwestern to Southeastern regions following already observed gradients of production and biodiversity (Coll et al., 2010; Mouillot et al., 2011). In OSMOSE-MED, the Western Mediterranean Sea accounted for 35 % of the total biomass, the Adriatic Sea 9 %, the Ionian and central Mediterranean Sea 31 % while the Aegean and Levantine Sea accounted for 26 % of the total biomass. The total biomass in the Adriatic Sea may be underestimated in view of the results found with a Mediterranean Ecopath model (Piroddi et al., 2015a). In this latter model, the Adriatic Sea was the area with the highest total biomass, followed by the Western Mediterranean Sea and the Ionian and Eastern Seas. This is partly due to the underestimation by the Eco3M-S biogeochemical model of the concentration in phytoplankton in this area (Kessouri, 2015).The Eastern basin appeared highly oligotrophic with low biomass values from OSMOSE-MED, with the exception of the Gulf of Gabès and waters surrounding the Nile plume, two regions that have been both characterized by high productivity (Hattab et al., 2013a).

The spatial distribution of catches, resulting from uniformly distributed fishing effort, globally followed the spatial distribution of the biomass with relatively less catches in the high sea (Figure 8). As for biomass, a low gradient of catch is predicted from the North to the South and from the West to the East, following the Mediterranean productivity pattern (Bosc et al., 2004; Ignatiades et al., 2009). The Iberian shelf waters, the Balearic Sea, the Gulf of Lions, the North Tyrrhenian Sea, the Adriatic Sea, the south Sicily, the Gulf of Gabès and the north Aegean Sea were all identified as hotspots of exploitation and concentrated most of the catches at the whole Mediterranean scale. Most of these areas were identified as highly impacted areas (Micheli et al., 2013a), in particular by demersal fishing activities and climate-induced changes, and coincide with the areas of conservation concern identified by Coll et al. (2012).

The analysis of the distribution of the mean size of the community revealed a clear gradient from the Northwestern to the Southeastern regions (Figure 9). Despite the fact that small pelagic fish species were mainly concentrated in the Northwestern region, the mean size weighted by abundance values was higher in the northern part of the basin. Some authors have argued that, the high salinity and temperature, the low productivity or a combination of all these factors were responsible for the “Levantine nanism phenomenon” that induces small body sizes for all species in general (Por, 1989; Sharir et al., 2011; Sonin et al., 2007). In OSMOSE, the growth in size is linked to the predation success. If the predation success is lower than a critical predation efficiency corresponding to maintenance requirements, fish can starve and growth rate is reduced (Shin and Cury, 2001). Thus, the very oligotrophic conditions of the eastern Mediterranean Sea could lead to reduced growth rates and smaller sizes for some species in OSMOSE-MED. The analysis of the spatial distribution of the mean size also suggested that large individuals were found in the Western high sea where catches were lower (Figure 9). On the one hand, large fish species abundances (e.g. *Thunnus thynnus* or *Xiphias gladius*) were more important in the Western high sea locally, which explains the large mean body size. On the other hand, the small sizes found in certain areas (e.g., Balearic Island, Northern Adriatic Sea, Cyprus Island) could be the result of heavy fishing, as fishing preferentially harvest larger-bodied individuals (from a given species, or species with larger mean size) but also induce a selection of slow growing individuals (Jørgensen et al., 2007; Law, 2000; Shin et al., 2005).

**Figure 9.**
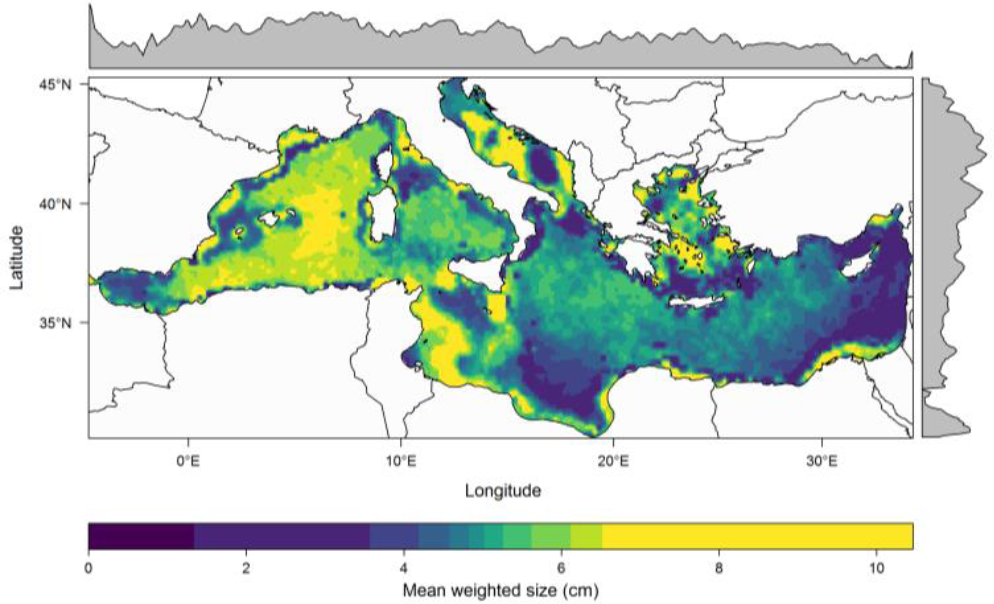
Spatial distribution of mean size (mean size weighted by species abundance) expressed in cm. Margins represent the meridionally and zonally averaged weighted size. (in color)

## 4 Conclusion and perspectives

### 4.1 The OSMOSE-MED challenge

In this paper, we described for the first time OSMOSE-MED, an integrated end-to-end model based on the coupling between a physical model (NEMOMED 12), a low trophic levels model (Eco3M-S) and a high trophic levels model (OSMOSE), representing the ecosystem dynamics and the trophic structure of the Mediterranean Sea in the 2006-2013 period. Numerous trophic modelling works have been realized in the Mediterranean Sea, most often at local scale (Bănaru et al., 2013; Coll et al., 2007; Corrales et al., 2017b, 2017a; Halouani et al., 2016; Hattab et al., 2013a) and rarely at the basin scale (Albouy et al., 2014; Piroddi et al., 2015a, 2017). The present study is the first attempt of an end-to-end trophic approach at the scale of the Mediterranean Sea, with an explicit modelling of the multispecies, spatial, life traits based and whole life cycle of the dynamics of a hundred interacting species. This was a challenge in many respects. The OSMOSE model, originally developed by Shin and Cury (2004, 2001), has never been tested for such a great number of species in interactions and at such a wide spatial scale. As noted by Fu et al. (2017), no more than 10 to 15 key species were usually included in an OSMOSE model. A few reasons could explain the selection of a restricted number of species to be modelled: (i) the extensive information and data required on species life histories to properly parameterize a model, (ii) the computation time and memory capacity to fit the model to observations, (iii) the desire to focus on major species and interactions only to simplify the complexity of the system. Here, we chose to move the modelling approach a step forward by being much more comprehensive in the explicit modelling of a large number of marine species in the Mediterranean Sea. Our ultimate goal was to build a tool representing the diversity of species and their interactions in a realistic way at the scale of the Mediterranean Sea, to be able to address the future impacts of climate change (e.g., species distribution shifts and plankton production changes) combined with other anthropogenic drivers on biodiversity such as fishing, its spatial dynamics across the whole Mediterranean basin and across Geographical Sub-Areas (GSAs), and the potential cascading effects on food webs and ecosystem services. We could clearly take advantage of existing historical collections of biological and ecological data that were assembled in various databases by the time of the project onset. In addition, the access to the recently upgraded High Performance Calculation (HPC) platform DATARMOR freed us from some technical barriers, and allowed us to envisage the calibration of a complex model such as OSMOSE-MED.

The first challenge, as for most end-to-end models, was to search and integrate a large amount of data and information from various sources, databases, scientific and “grey” literatures as well as outputs from other models (de Mora et al., 2016; Fulton, 2010). To our knowledge, OSMOSE-MED is the most complete model built at the whole Mediterranean Sea scale, in terms of species and process representativeness. It integrates the best ecological knowledge in the Mediterranean Sea despite some gaps can be pointed out, mostly concerning for the fish species in the southern part of the basin (Dimarchopoulou et al., 2017). There is, for instance, no biological information for as many as 43 % of the Mediterranean fish species (Dimarchopoulou et al., 2017). The lack of biological and ecological data for a large number of species as well as the quality of commercial fisheries data, especially in the southern and eastern parts of the Mediterranean Sea, are hindrances to reliable stock assessments, to the development of more integrated ecosystem models, and thus to the implementation of an effective ecosystem-based management to achieve good environmental status in the Mediterranean Sea (Coll et al., 2013; Piroddi et al., 2015a, 2017). A crucial challenge is to increase the number of assessed stocks in order to ensure their sustainable exploitation in a first step, to allow the parameterization and calibration of more integrated ecosystem models that would support the development of ecosystem-based fisheries management at the Mediterranean basin scale in a second (Cardinale and Scarcella, 2017; Coll et al., 2013; Colloca et al., 2013). Moreover, the region is generally suffering from the problem of data ownership, reliability, and accessibility (Katsanevakis et al., 2015).

The second challenge was to maintain in co-existence all HTL species and provide a realistic representation of the biodiversity. It was probably the most critical and time consuming step, given the stochasticity and the complexity of the model. The number of trophic links, the connectance, and the importance of feedback controls can be large in an OSMOSE model and can render the calibration procedure complicated and fastidious (Halouani et al., 2016; Marzloff et al., 2009; Travers-Trolet et al., 2014). We exploited the capacities of the evolutionary optimization algorithm in order to find a set of estimated parameters within a 195-dimensional search space (Oliveros-Ramos et al., 2017; Oliveros-Ramos and Shin, 2016) that reproduced state variables and indicators close to observations. The calibrar R package was used for the first time for such a complex model configuration (large number of parameters, stochastic model with many nonlinearities) and proved its capacity to solve complicated minimization problems (Oliveros-Ramos and Shin, 2016). For computational time reasons, and the need for continuous iterative trials and feedbacks between model’s parameterization and observations, the calibration of OSMOSE-MED has taken more than one year and required the use of high performance computing facilities. The development of OSMOSE-MED represents a significant advance for OSMOSE and calibrar user’s communities, but also more broadly in the field of ecosystem modelling, as a proof of concept that complex representation of species dynamics and their interactions can be achieved and can produce realistic spatial, multispecies, and whole life cycle dynamics under the influence of climate and anthropogenic drivers.

### 4.2 Limitations of the model

Ecosystem models, despite their increasing complexity, granularity and representativeness are always idealised or simplified conceptual representations of very complex systems (Gunawardena, 2014). As all models involve some simplifications, certain limitations exist and can be discussed:

#### Benthos compartment

an important bentho-pelagic coupling exists in the Mediterranean Sea and has been highlighted in several Ecopath models in the region (Banaru et al., 2013; Coll et al., 2007; Corrales et al., 2015; Hattab et al., 2013a). Moreover, many species included in OSMOSE-MED have omnivorous and carnivorous diets partly based on benthic organisms such as polychaetes, amphipods or crustaceans. This is why a benthos “black box” has been added in OSMOSE-MED, with a constant biomass and a uniform spatial distribution. Given the importance of this trophic compartment in the Mediterranean Sea, this compartment would deserve an improved representation, for example by considering multiple functional groups having common biological and ecological characteristics (e.g. meiofauna, bivalves, echinoderms) (Grüss et al., 2016). As these new developments are clearly impeded by the lack of data for both parameterization and calibration of the model, an intermediate complexity approach could be adopted by modelling these more refined benthic compartments as “background taxa” for which only predation, mean growth rate and spatial distribution are modelled. This new category of species of intermediate complexity, allowing to cope with limited datasets and to include more species of interest while keeping the model reasonably complex, was recently coded in OSMOSE (Fu et al., 2017).

#### Ontogenic changes in habitats

Numerous species included in OSMOSE-MED exhibit clear ontogenic shifts in habitats in the Mediterranean Sea (Cartes et al., 2009; Druon et al., 2015, 2016; Giannoulaki et al., 2013b, 2013a; Macpherson, 1998). These ontogenic range shifts can play a critical role in population dynamics and ecosystem functioning (MacCall, 1990; Macpherson and Duarte, 1991; Methratta and Link, 2007). For instance, Caddy (1990) hypothesized that the sustainability of the majority of Mediterranean fisheries depended on spawners refuging on continental slopes. For most of the major commercial species (including hake, monkfish and shrimps) the continental slope and canyons are used as spawning areas that are less accessible to fishing fleets, while the continental shelf and the strip coast that are more intensively fished, are preferred zones for nurseries (Würtz, 2012). Therefore, including different spatial distribution maps (i.e. spawning and nursery grounds) for some key species like small pelagic fish (e.g. European anchovy, European pilchard or European mackerel) and demersal fish (e.g. European hake, Red mullet) could potentially improve the spatial representation of food webs and population dynamics, and their vulnerability to fishing. Habitat suitability models by stage or size class, that relate abundance information from surveys with environmental variables could be used in this purpose (Druon et al., 2015; Giannoulaki et al., 2013a).

#### Spatialized fishing effort/mortality

As discussed above, OSMOSE-MED considered a uniform spatial distribution of fishing effort. Fishing effort being mainly distributed along the coasts and on the continental shelf, this assumption is not realistic (Kroodsma et al., 2018; Leleu et al., 2014; Maynou et al., 2011; Ramírez et al., 2018), though the lower biomass in the open sea enables to counterbalance this potential source of bias (Figure 8). Moreover, most fisheries targeting large pelagic fish such as tunas or swordfish operate in the open sea, due to the target species distribution pattern (Druon et al., 2016). Fishing effort metadata, reported at the species and Geographical-Sub area scales, from the Data Collection Reference Framework (GFCM, 2018) could be available in order to improve the differential pressures exerted by fishing across the Mediterranean sea. Another option to spatialize fishing effort/mortality would be to model as many exploited populations in a species as the number of evaluated stocks. This supposes to know the true number of stocks in the Mediterranean Sea and the possible connectivities between them (Fiorentino et al., 2014; Ragonese et al., 2016).

#### Uncertainty

Marine ecosystems are structurally complex, spatially and temporally variable, difficult and costly to observe, all of which can potentially lead to considerable uncertainty in model predictions(Cheung et al., 2016; Hill et al., 2007; Payne et al., 2016). There are many sources of uncertainty in ecosystem models, from the structural (model) uncertainty and the initialization and internal variability uncertainty to the parametric uncertainty (Payne et al., 2016). Assessing these different types of uncertainty would allow to build confidence intervals around our predictions with the OSMOSE-MED model and would clearly increase the relevance of using it in projections and in support of decision-making in the Mediterranean Sea (Gal et al., 2014; Hill et al., 2007; Hyder et al., 2015; Payne et al., 2016).

In the present study, the uncertainty due to the sources of input data (i.e. parametric uncertainty) could be tested as a first step. Most of the data used for parameterizing OSMOSE-MED come from the modelled area but some parameters for data-poor species (e.g., relative fecundity, growth parameters) were obtained from ecosystems outside the Mediterranean region and can differ considerably according to the ecosystem (Halouani et al., 2016). Sensitivity analysis on such parameters could be tested following the methodology employed in Lehuta et al. (2010) or Ortega-Cisneros et al. (2017).

### 4.3 Potential uses of OSMOSE-MED

The OSMOSE-MED model is an integrated ecosystem model addressing the combined effects of fishing and climate change on marine biodiversity at the whole Mediterranean basin scale to provide scientific support to strategize fisheries management.

The model can for example provide insights on the impacts of climate change on operational fisheries reference levels, such as Maximum Sustainable Yield (MSY) and multi-species MSY at the Mediterranean Sea scale (Lehuta et al., 2016). It has also the capacity to guide the prioritization of multiple spatial conservation plans such as the implementation of coherent Marine protected Areas (MPAs) networks by European states (Lehuta et al., 2016; Liquete et al., 2016; Micheli et al., 2013b) as required by the Marine Strategy Framework Directive (MSFD) (European Commission, 2008). Moreover, numerous biodiversity and food webs MSFD indicators can be directly derived from outputs of OSMOSE-MED, making this model a relevant tool to help the planning and integration of policies like the MSFD that seeks to achieve, for all European seas, Good Environmental Status, by 2020 (Cardoso et al., 2010; Piroddi et al., 2015b). Model outputs can be used to provide an evidence base to inform decision-making, especially in the frame of the EU’s Blue growth strategy that supports sustainable growth in the marine and maritime sectors as a whole (European Commission, 2017), and the General Fisheries Commission for the Mediterranean (GFCM) mid-term strategy (2017-2020) that has been developed to support the achievement of the United Nations targets (e.g., the Sustainable Development Goal 14) (GFCM, 2017b)

OSMOSE-MED model can also be viewed as a tool to communicate effectively with managers and other non-scientist end users of Mediterranean ecosystems and help incorporating scientific evidence into environmental decision-making (Cartwright et al., 2016; Jönsson et al., 2015; Rose et al., 2010).

## Acknowledgements

The authors acknowledge the Pôle de Calcul et de Données Marines (PCDM) for providing DATARMOR computational resources. URL: http://www.ifremer.fr/pcdm. We are grateful to the participants to the MEDITS survey program which has been conducted within the Data Collection Framework (DCF) since 1994. We also thank Sabrine Drira for her help in developing species distribution models. Fabien Moullec was funded by a PhD grant from the French Ministry of higher Education, Research and Innovation. This work was partially funded by the *USBIO* project of the LabEx CeMEB, an ANR “Investissements d’avenir” program (ANR-10-LABX-04-01).

## Author contributions

F.M. developed the model, acquired the data, analyzed and interpreted the data. L.V., P.V., N.B., F.G. and Y-J.S. helped in developing the model. Y-J.S. helped in data analysis and interpretation. C.U. provided data on primary and secondary productions (from the biogeochemical model). P.V. and N.B. helped with the programming code of OSMOSE and use of the HPC cluster DATARMOR. P.C., A.E., C.F., M.G., A.J., A.L., E.L.D., P.M., P.P., M.T.S., I.T. and M.V. provided data from the MEDITS survey. F.M. led the drafting of the manuscript with the contributions and revisions from all the authors.

