## Supplementary materials for "Catching the big picture of the Mediterranean Sea biodiversity with an end-to-end model of climate and fishing impacts"

#### Appendix A. Details and equations of OSMOSE model

The OSMOSE 3 update 2 used in this paper is freely available on the OSMOSE website (www.osmose-model.org). The OSMOSE model aims at exploring fish community dynamics and the ecosystem effects of fishing and climate change. This Individual-Based Model (IBM) assumes opportunistic predation based on spatio-temporal co-occurrence and size adequacy between a predator and its prey. Individuals are grouped in schools which are characterized by their size, weight, age, taxonomy and geographical location on the 2D grid. At each time step, the main processes of marine species life cycle occur:

##### 1- Spatial distribution

The spatial distribution of super individuals/schools at each time step is driven by input maps that are species dependent (ontogenic or seasonal changes of spatial distribution were not considered in this study). At each time step $t$, when new eggs are released, schools are uniformly distributed over their specific distribution area.
As the maps do not change from one-time step to the next, schools can move to adjacent cells within their distribution area following a random walk process. Range of the random walk is expressed in number of cells. If range = zero, the school remains in the current cell. If range = 1, the school can either stay in the current cell or move in any of the 8 immediately adjacent cells. If range=2, the school can either stay in the current cell or move in any of the 24 immediately adjacent cells. Random walk movements are meant to represent small-scale foraging movement and diffusion.

##### 2- Mortalities

By default, the OSMOSE version used to build OSMOSE-MED use a stochastic algorithm to implement the different mortality sources: within each time step, the total mortality $Z$ of a given school $j$ ($Z_j$) is comprised of predation mortality caused by various schools $\{i\}$ ($\sum M_{predation\ j,\ i}$), starvation mortality ($M_{starvation\ j}$), fishing mortality ($F_j$) and other natural mortality sources not explicitly represented in the model ($M_{nat\ j}$) (e.g. disease, predation mortality due to predators not included in the model).

###### a. *Predation mortality*

Super individuals/schools interact locally through predation events in a stochastic manner. In OSMOSE, predation is an opportunistic process based on the spatial overlap between predators and potential prey and size adequacy between the predator and the potential prey (Shin and Cury, 2004, 2001). Predator/prey size ratios are defined by species, and age/stage in case of ontogenic changes in feeding behaviour (e.g., shift from particulate feeding to filter feeding). The amount of prey eaten depends on the local relative biomass of prey, the biomass of potential competitors, and on the maximum food edible by the predator (Travers et al., 2009). An accessibility matrix (between predators and prey), which can depend on the vertical distribution of each species, is also defined. This matrix can also be used to restrict the range of possible prey for more selective predators as is the case for benthic organisms compared to pelagic ones.
The total accessible biomass of prey to school $i$ in a cell $(x, y)$ at time step $t$ $(AP_{i,x,y,t})$ is calculated following equation:

$$AP_{i,x,y,t} = \sum_j a_{i,j} \,.\, B_{j,x,y,t} \quad | \quad \frac{L_i}{sr_{max}} < L_j < \frac{L_i}{sr_{min}} \qquad \text{Equation A.1.}$$

$$\begin{cases} if\ AP_{i,x,y,t} > r_i \,.\, B_{i,x,y,t} & PB_{i,j,\Delta t} = a_{i,j} \,.\, B_{j,t} \,.\, \frac{r\,.\,B_{i,t}}{AP_{i,x,y,t}} \\ if\ AP_{i,x,y,t} \leq r_i \,.\, B_{i,x,y,t} & PB_{i,j,\Delta t} = a_{i,j} \,.\, B_{j,t} \end{cases} \qquad \text{Equation A.2.}$$

Where $L$ is the length of prey school $j$ or predator school $i$, $B_{j,x,y,t}$ is the biomass of prey school $j$ in the cell $(x, y)$ at time step $t$, $a_{i,j}$ is the accessibility coefficient of prey $j$ to predator $i$, $sr$ is the suitable predator-prey size ratio, $\mathrm{r_i}$ is the maximum ingestion rate of species $i$, and $\mathrm{PB_{i,j,\Delta t}}$ the biomass of $j$ preyed upon by school $i$ during a time step $t$.
A predation efficiency is calculated for each school $i$ $(\xi_i)$. This coefficient is determined by the ratio between the food biomass ingested by a group per time step and the maximum food ingestion $(\mathrm{r_i} * Bi, x, y, t)$. Depending on the predation efficiency, schools can grow or starve.

*b. Starvation mortality*

When predation efficiency $\xi_{i,t}$ is below the critical value $\xi_{crit}$ (corresponding to maintenance requirements), schools $i$ undergo a starvation mortality which increases linearly with the decrease of predation efficiency $\xi_{i,t}$ and leads to a decrease of the school abundance (Shin and Cury, 2004, 2001):

$$M_{starvation\ i,t} = M_{starvation\ i}^{max} - \frac{M_{starvation\ i}^{max}}{\xi_i^{crit}}\ \xi_{i,t} \qquad \text{Equation A.3.}$$

c. *Fishing mortality*

Fishing mortality $(F_{i,t})$ is applied to exploited schools whose individuals are larger than a defined size or older than a defined age at recruitment. $F_{i,t}$ is homogeneous spatially but can vary over time following a fishing seasonality provided as input for each species (Travers et al., 2009).

*d. Natural mortality*

An additional source of natural mortality (i.e. mortality due to marine organisms and events that are not explicitly considered in OSMOSE) is applied to schools older than 1 month.

The mortality of eggs and larvae $M_0$ applied to 0-1 month old individuals accounts for the high mortality undergone by early life stages and critical stages such as first feeding larvae (Travers et al., 2009). This parameter is typically estimated through the calibration of the model to observations.

##### 3- Growth

Mean growth rate in length of a school $i$ of species $s$ and of age $a$ are calculated from the von Bertalanffy model. The variability around the mean depends on food ingestion at each time step. The von Bertalanffy growth model only applies for schools older than a threshold age ($A_{thres}$) defined from the literature. Below that threshold, growth is assumed to be linear. Assuming a linear growth between age 0 day and $A_{thres}$ ensures a more realistic computation of mean length increases for early ages of HTL species (Grüss et al., 2015; Travers et al., 2009).

$$\Delta L_{i,s,a} = L_{\infty s}\,(1 - e^{-K_s})e^{-K_s(a-a_{0s})}$$ *Equation A.4.*

The actual growth rate of a fish takes into account the quantity of food ingested by each school $i$ during a time step $t$: individuals of school $i$ grow in size and weight when their predation efficiency at $t$ is greater than $\xi_{crit}$. The threshold $\xi_{crit}$ corresponding to basic maintenance requirements is set by default for all species at 0.57 (Laevastu and Larkins, 1981). For a school $i$, if $\xi_i \geq \xi_{crit}$, growth rate in length varies linearly with $\xi_i$ such that the median value between $\xi_{crit}$ and 1 ($\xi_{max}$) corresponds to the mean von Bertalanffy growth rate Thus, the growth rate in length of a school $i$, of age $a$, of species $s$, and at time $t$, follows the expression:

$$\begin{cases} \Delta L_{s,a,i,t} = 0 & if\ \xi_{i,t} < \xi_{crit} \\ \Delta L_{s,a,i,t} = \frac{2\Delta L_{s,a,}}{1-\xi_{crit}}\,(\xi_{i,t} - \xi_{crit}) & if\ \xi_{i,t} \geq \xi_{crit} \end{cases}$$ *Equation A.5.*

The mean body weight of a school (or cohort) $i$ of species $s$ at time $t$ is subsequently calculated from the allometric relationship:

$$W_{i,s,t} = c\,L_{i,s,t}^{b}$$ *Equation A.6.*

Where the $c$ parameter is the condition factor and $b$ the allometric power.

##### 4- Reproduction

At the end of each time step, the spawning stock biomass of a species is calculated (biomass of all fish which length is greater than the length at sexual maturity $L_{mat}$). The numbers of eggs spawned by a species $s$ at time $t$ ($N_{s,0,t}$) is calculated as follows:

$$N_{s,0,t} = SR_s * \phi_s * \Theta^{time\ step} * (SSB_{s,t})$$ *Equation A.7.*

With $SR_s$, $\phi_s$, $\Theta^{\text{time step}}$and $SSB_{s,t}$ representing the female:male sex ratio of the species $s$, the relative fecundity (number of eggs spawned per gram of mature female per year), the probability for species $s$ to spawn within a given time step (spawning seasonality), the spawning stock biomass of species $s$ at time $t$, respectively.

As growth variability is implemented in relation to food intake, the reproductive success also depends implicitly on the food conditions that are encountered, locally in time and space, by each school (Shin and Cury, 2004).

#### Appendix B. Input parameters of OSMOSE-MED

Table B.1. Input parameters of the OSMOSE-MED for each of the 100 species modelled. *$L_\infty$, K, $t_0$*: the von Bertalanffy growth parameters; *b*: the exponent of the allometric length-weight relationship, *c*: constant of proportionality of the allometric length-weight relationship; Lmat: size at maturity; Θ: relative fecundity; Amax: longevity; D: Mortality rate due to predation from other species that are not explicitly considered in the model; Lrec: size at recruitment; Arec: age at recruitment. (Lpred / Lprey): minimum and maximum predation ratios for juveniles and adults. For the same parameter, if different estimates were available in the Mediterranean Sea, the average was considered. See Appendix C for the corresponding list of references.

| Common name | Scientific name | Growth | | | | | Reproduction | | Mortality | | | | | | Diet | | | | | |
|---|---|---|---|---|---|---|---|---|---|---|---|---|---|---|---|---|---|---|---|---|
| | | $L_\infty$ | $K$ | $t_0$ | $c$ | $b$ | Θ | $L_{mat}$ | $D$ | $A_{max}$ | $L_{rec}$ | $A_{rec}$ | Mean catch | (Lpred / Lprey) max | | | (Lpred / Lprey) min | | | $L_{thres}$ |
| | | (cm) | ($yr^{-1}$) | (yr) | ($g\ cm^{-b}$) | | (eggs $g^{-1}.year^{-1}$) | (cm) | ($yr^{-1}$) | (yr) | (cm) | (yr) | (t) | juvenile | | adult | juvenile | | adult | (cm TL) |
| Allis shad | *Alosa alosa* | 70.300 | 0.351 | 0.000 | 0.00359 | 3.21267 | 140 | 40.0 | 0.24 | 8 | null | 1 | 1605 | 35 | ; | 35 | 70 | ; | 70 | null |
| Twaite shad | *Alosa fallax* | 52.100 | 0.241 | -0.580 | 0.00548 | 3.28350 | 143 | 22.5 | 0.20 | 8 | null | 1 | 1811 | 5 | ; | 5 | 30 | ; | 30 | null |
| European eel | *Anguilla anguilla* | 70.187 | 0.408 | -0.502 | 0.00125 | 3.18882 | 3906 | 60.0 | 0.26 | 13.2 | null | 1 | 1053 | 8 | ; | 8 | 25 | ; | 25 | null |
| Meagre | *Argyrosomus regius* | 140.000 | 0.145 | -0.225 | 0.01739 | 2.85120 | 900 | 53.3 | 0.28 | 42 | null | 1 | 1988 | 6 | ; | 6 | 85 | ; | 85 | null |
| Giant red shrimp | *Aristaeomorpha foliacea* | 7.139 | 0.444 | -0.140 | 0.00135 | 2.63000 | 5477 | 3.7 | 0.72 | 8 | 2.5 | null | 65 | 4.5 | ; | 7.5 | 50 | ; | 50 | 3 |
| Blue and red shrimp | *Aristeus antennatus* | 7.433 | 0.331 | -0.078 | 0.28801 | 2.46267 | 9386 | 2.6 | 0.90 | 10 | 2.5 | null | 5802 | 4.5 | ; | 7.5 | 50 | ; | 50 | 3 |
| Big-scale sand smelt | *Atherina boyeri* | 14.750 | 0.210 | -0.925 | 0.00677 | 3.09347 | 784 | 5.4 | 0.15 | 4 | null | 1 | 420 | 15 | ; | 15 | 100 | ; | 100 | null |
| Bullet tuna | *Auxis rochei rochei* | 57.388 | 0.181 | -4.155 | 0.05420 | 2.68000 | 242 | 35.8 | 0.21 | 5 | null | 1 | 6753 | 5 | ; | 5 | 20 | ; | 20 | null |
| Garfish | *Belone belone* | 54.800 | 0.351 | -1.463 | 0.00173 | 2.88150 | 11 | 30.6 | 0.20 | 8 | null | 1 | 1571 | 8 | ; | 8 | 82 | ; | 82 | null |
| Bogue | *Boops boops* | 32.400 | 0.180 | -1.363 | 0.00676 | 3.13503 | 1396 | 13.0 | 0.40 | 5 | 13 | null | 36904 | 15 | ; | 15 | 150 | ; | 150 | null |
| Blue runner | *Caranx crysos* | 41.200 | 0.350 | 0.000 | 0.00890 | 3.11000 | 790 | 23.1 | 0.30 | 11 | null | 1 | 0 | 2 | ; | 2 | 10 | ; | 10 | null |
| Tub gurnard | *Chelidonichthys lucerna* | 55.057 | 0.416 | -0.467 | 0.01197 | 2.95491 | 587 | 20.2 | 0.29 | 14.8 | 21.5 | null | 540 | 6 | ; | 6 | 85 | ; | 85 | null |
| Mediterranean rainbow wrasse | *Coris julis* | 27.200 | 0.146 | -1.513 | 0.00788 | 3.14100 | 78 | 16.2 | 0.20 | 6 | null | 1 | 53 | 6 | ; | 6 | 350 | ; | 350 | null |
| Common dolphin fish | *Coryphaena hippurus* | 101.450 | 1.659 | 0.036 | 0.00970 | 3.01370 | 120 | 68.4 | 0.30 | 3.66 | 27.8 | null | 4866 | 2.58 | ; | 2.58 | 10 | ; | 10 | null |
| Common shrimp | *Crangon crangon* | 7.790 | 1.170 | 0.000 | 0.01020 | 2.80600 | 815 | 2.5 | 0.55 | 2 | null | 1 | 631 | 2.5 | ; | 3 | 200 | ; | 200 | 0.35 |
| Cristal goby | *Crystallogobius linearis* | 5.400 | 0.970 | -0.300 | 0.00589 | 3.13000 | 8050 | 2.7 | 0.20 | 1 | null | 1 | 0 | 4 | ; | 4 | 20 | ; | 20 | null |
| Common dentex | *Dentex dentex* | 90.625 | 0.088 | -2.362 | 0.01150 | 3.12029 | 650 | 38.6 | 0.23 | 26 | null | 1 | 3705 | 6 | ; | 6 | 85 | ; | 85 | null |
| Pink dentex | *Dentex gibbosus* | 107.240 | 0.120 | -0.900 | 0.00820 | 3.13000 | 311 | 41.5 | 0.27 | 16 | null | 1 | 47 | 6 | ; | 6 | 85 | ; | 85 | null |
| Morocco dentex | *Dentex maroccanus* | 34.900 | 0.170 | -1.720 | 0.06370 | 2.73500 | 448 | 10.0 | 0.25 | 10 | null | 1 | 0 | 6 | ; | 6 | 35 | ; | 35 | null |
| European seabass | *Dicentrarchus labrax* | 69.620 | 0.247 | -0.666 | 0.01054 | 3.07277 | 417 | 26.5 | 0.10 | 7 | 22.8 | null | 6983 | 8 | ; | 8 | 60 | ; | 60 | null |
| Annular sea bream | *Diplodus annularis* | 22.252 | 0.281 | -1.229 | 0.01501 | 3.12911 | 400 | 10.6 | 0.20 | 13 | 10 | null | 4969 | 6 | ; | 6 | 20 | ; | 20 | null |
| Zebra seabream | *Diplodus cervinus* | 68.800 | 0.110 | -0.750 | 0.01160 | 3.14000 | 199 | 25.0 | 0.20 | 17 | 10 | null | 1 | 11 | ; | 11 | 20 | ; | 20 | null |
| Sharpsnout seabream | *Diplodus puntazzo* | 45.280 | 0.191 | -0.306 | 0.02450 | 3.00897 | 3233 | 21.9 | 0.20 | 18 | 10 | null | 108 | 7 | ; | 7 | 20 | ; | 20 | null |

| | | | | | | | | | | | | | | | | | | | | |
|---|---|---|---|---|---|---|---|---|---|---|---|---|---|---|---|---|---|---|---|---|
| White sea bream | *Diplodus sargus sargus* | 44.200 | 0.152 | -1.483 | 0.01715 | 3.08188 | 200 | 20.8 | 0.20 | 10 | 15 | null | 5923 | 6 | ; | 6 | 20 | ; | 20 | null |
| Common two-banded sea bream | *Diplodus vulgaris* | 30.018 | 0.195 | -1.728 | 0.01861 | 3.10391 | 162 | 17.4 | 0.10 | 7 | 12.5 | null | 3683 | 7 | ; | 7 | 20 | ; | 20 | null |
| Horned octopus | *Eledone cirrhosa* | 19.280 | 0.387 | -0.030 | 0.59682 | 2.62700 | 7 | 7.8 | 0.10 | 2 | 3.75 | null | 7126 | 5 | ; | 5 | 25 | ; | 25 | null |
| European anchovy | *Engraulis encrasicolus* | 18.408 | 0.473 | -1.745 | 0.00527 | 3.16273 | 414 | 10.5 | 0.10 | 4.5 | 10 | null | 136759 | 5 | ; | 10 | 500 | ; | 700 | 8 |
| White grouper | *Epinephelus aeneus* | 144.325 | 0.166 | -0.767 | 0.01090 | 3.01500 | 515 | 55.0 | 0.26 | 17 | null | 1 | 256 | 5.5 | ; | 5.5 | 40 | ; | 30 | 50 |
| Dusky grouper | *Epinephelus marginatus* | 137.467 | 0.090 | -0.767 | 0.01230 | 3.05000 | 334 | 65.0 | 0.10 | 50 | null | 1 | 1530 | 4.5 | ; | 4.5 | 40 | ; | 30 | 50 |
| Red-eye round herring | *Etrumeus teres* | 33.770 | 0.200 | -1.631 | 0.00900 | 3.12159 | 196 | 13.7 | 0.20 | 3 | null | 1 | 2 | 10 | ; | 10 | 500 | ; | 500 | 10 |
| Grey gurnard | *Eutrigla gurnardus* | 46.000 | 0.160 | 0.000 | 0.00588 | 3.14714 | 404 | 21.0 | 0.32 | 21 | 12.5 | null | 1658 | 4 | ; | 4 | 100 | ; | 100 | null |
| Black-mouthed dogfish | *Galeus melastomus* | 64.000 | 0.150 | 0.000 | 0.00250 | 3.02000 | 193 | 44.5 | 0.35 | 8 | 28.5 | null | 382 | 4 | ; | 4 | 14 | ; | 14 | null |
| Black goby | *Gobius niger* | 15.900 | 0.391 | -1.328 | 0.01241 | 3.00141 | 1563 | 7.8 | 0.20 | 4.33 | null | 1 | 182 | 5 | ; | 5 | 26 | ; | 26 | null |
| Lusitanian toadfish | *Halobatrachus didactylus* | 52.000 | 0.130 | 0.000 | 0.02022 | 3.02375 | 9 | 28.2 | 0.05 | 12 | null | 1 | 2 | 1.1 | ; | 1.1 | 5.82 | ; | 5.82 | null |
| Shortfin squid | *Illex coindetii* | 29.630 | 2.000 | -0.096 | 0.01880 | 3.20000 | 1778 | 14.1 | 0.30 | 1.13 | 6.25 | null | 3620 | 5 | ; | 5 | 40 | ; | 40 | null |
| Megrim | *Lepidorhombus whiffiagonis* | 44.800 | 0.170 | -1.070 | 0.00640 | 2.99300 | 603 | 22.4 | 0.33 | 7 | 10 | null | 178 | 2 | ; | 6 | 15 | ; | 13 | 15 |
| Golden grey mullet | *Liza aurata* | 43.214 | 0.247 | -0.467 | 0.01262 | 2.90264 | 1152 | 24.7 | 0.53 | 8 | null | 1 | 759 | 15 | ; | 15 | 100 | ; | 100 | null |
| Thinlip grey mullet | *Liza ramada* | 44.335 | 0.288 | -0.411 | 0.01228 | 2.95890 | 553 | 28.3 | 0.42 | 10 | null | 1 | 0 | 40 | ; | 40 | 100 | ; | 100 | null |
| Leaping mullet | *Liza saliens* | 38.545 | 0.229 | -0.359 | 0.01750 | 2.91513 | 1822 | 21.9 | 0.34 | 4 | null | 1 | 0 | 40 | ; | 40 | 100 | ; | 100 | null |
| European squid | *Loligo vulgaris* | 23.800 | 1.740 | 0.060 | 0.05000 | 2.41810 | 211 | 13.8 | 0.40 | 1.25 | 3.25 | null | 96 | 10.3 | ; | 10.3 | 116.7 | ; | 116.7 | null |
| Black-bellied angler | *Lophius budegassa* | 103.000 | 0.150 | -0.050 | 0.02440 | 2.84570 | 102 | 33.5 | 0.37 | 7 | 6.25 | null | 121 | 4.1 | ; | 4.1 | 10.82 | ; | 10.82 | null |
| Anglerfish | *Lophius piscatorius* | 102.000 | 0.150 | -0.050 | 0.02113 | 2.88043 | 102 | 34.0 | 0.27 | 8 | 9 | null | 3608 | 4.2 | ; | 4.2 | 7.73 | ; | 7.73 | null |
| Whiting | *Merlangius merlangus* | 39.748 | 0.143 | -1.623 | 0.01266 | 3.05593 | 1700 | 24.5 | 0.24 | 13 | 7.5 | null | 4544 | 1.8 | ; | 1.8 | 6 | ; | 6 | null |
| European hake | *Merluccius merluccius* | 105.423 | 0.189 | -0.049 | 0.00434 | 3.09755 | 203 | 28.9 | 0.10 | 5.5 | 7.5 | null | 26862 | 1.5 | ; | 2 | 15 | ; | 12 | 15 |
| Blue whiting | *Micromesistius poutassou* | 49.300 | 0.297 | 0.000 | 0.00290 | 3.33267 | 120 | 21.0 | 0.20 | 8.5 | 12.5 | null | 9301 | 2 | ; | 2 | 25 | ; | 25 | null |
| Flathead grey mullet | *Mugil cephalus* | 63.700 | 0.315 | -0.275 | 0.00886 | 3.14745 | 1680 | 44.8 | 0.31 | 11 | null | 1 | 2582 | 35 | ; | 35 | 100 | ; | 100 | null |
| Red mullet | *Mullus barbatus barbatus* | 29.205 | 0.380 | -0.443 | 0.00771 | 3.12421 | 341 | 11.2 | 0.10 | 5 | 9 | null | 19749 | 6 | ; | 6 | 20 | ; | 20 | null |
| Striped red mullet | *Mullus surmuletus* | 36.023 | 0.290 | -1.279 | 0.00917 | 3.10120 | 786 | 15.5 | 0.21 | 6.2 | 10 | null | 10817 | 6 | ; | 6 | 16 | ; | 16 | null |
| Smooth hound | *Mustelus mustelus* | 175.000 | 0.090 | -2.845 | 0.00191 | 3.20650 | 0 | 107 | 0.01 | 24 | 50 | null | 367 | 8 | ; | 8 | 20 | ; | 20 | null |
| Norway lobster | *Nephrops norvegicus* | 22.700 | 0.324 | -0.290 | 0.00043 | 3.17278 | 68 | 9.5 | 0.20 | 15 | 1.25 | null | 6425 | 10 | ; | 10 | 50 | ; | 50 | null |
| Common octopus | *Octopus vulgaris* | 29.600 | 0.560 | -0.230 | 0.42000 | 2.98700 | 95 | 14.5 | 0.15 | 1.3 | 4 | null | 16796 | 5 | ; | 5 | 20 | ; | 20 | null |
| Axillary seabream | *Pagellus acarne* | 28.324 | 0.287 | -1.707 | 0.00972 | 3.19022 | 1134 | 18.0 | 0.43 | 8 | 12 | null | 2284 | 5 | ; | 5 | 25 | ; | 25 | null |
| Common pandora | *Pagellus erythrinus* | 37.351 | 0.188 | -1.147 | 0.02173 | 2.94847 | 156 | 16.4 | 0.24 | 6.6 | 11 | null | 11571 | 5 | ; | 5 | 25 | ; | 25 | null |
| Common seabream | *Pagrus pagrus* | 63.960 | 0.144 | 0.000 | 0.02217 | 3.03114 | 517 | 24.7 | 0.29 | 12 | 12 | null | 5743 | 6 | ; | 6 | 60 | ; | 60 | null |

| | | | | | | | | | | | | | | | | | | | | |
|---|---|---|---|---|---|---|---|---|---|---|---|---|---|---|---|---|---|---|---|---|
| Common prawn | *Palaemon serratus* | 8.596 | 0.605 | 0.000 | 0.00060 | 1.96950 | 1500 | 5.8 | 0.20 | 3 | null | 1 | 17 | 6 | ; | 7.5 | 700 | ; | 242 | 3 |
| Common spiny lobster | *Palinurus elephas* | 17.850 | 0.190 | -0.260 | 0.00450 | 2.62000 | 116 | 8.1 | 0.10 | 15 | 0.4 | null | 333 | 8 | ; | 8 | 100 | ; | 100 | 3 |
| Deep-water rose shrimp | *Parapenaeus longirostris* | 4.439 | 0.539 | -0.129 | 0.18924 | 2.49554 | 7486 | 2.2 | 0.40 | 6 | 1.1 | null | 18449 | 8 | ; | 12 | 100 | ; | 242 | 2 |
| Caramote prawn | *Penaeus kerathurus* | 18.030 | 0.690 | -0.940 | 0.00469 | 2.40600 | 7705 | 3.0 | 0.44 | 3 | 7 | null | 8325 | 10 | ; | 10 | 75 | ; | 75 | null |
| Forkbeard | *Phycis phycis* | 67.150 | 0.195 | -0.515 | 0.00520 | 3.18800 | 1833 | 32.2 | 0.42 | 5 | null | 1 | 850 | 5 | ; | 5 | 26 | ; | 26 | null |
| European flounder | *Platichthys flesus flesus* | 38.500 | 0.420 | -2.222 | 0.00677 | 3.10100 | 2611 | 25.0 | 0.19 | 15 | null | 1 | 243 | 22 | ; | 22 | 330 | ; | 330 | null |
| European plaice | *Pleuronectes platessa* | 55.635 | 0.146 | -1.504 | 0.00968 | 3.03033 | 206 | 23.9 | 0.13 | 32.5 | null | 1 | 0 | 25 | ; | 25 | 100 | ; | 100 | null |
| Bluefish | *Pomatomus saltatrix* | 111.201 | 0.173 | -0.904 | 0.01394 | 2.92726 | 97 | 37.0 | 0.35 | 9 | null | 1 | 8294 | 7 | ; | 7 | 50 | ; | 50 | null |
| Marbled goby | *Pomatoschistus marmoratus* | 6.453 | 1.500 | 0.000 | 0.00751 | 3.19933 | 1925 | 3.3 | 0.20 | 2 | null | 1 | 0 | 5 | ; | 5 | 15 | ; | 15 | null |
| Sand goby | *Pomatoschistus minutus* | 8.150 | 1.154 | -0.165 | 0.00370 | 3.28900 | 1229 | 4.0 | 0.20 | 2.6 | null | 1 | 0 | 5 | ; | 5 | 15 | ; | 15 | null |
| Common guitarfish | *Rhinobatos rhinobatos* | 128.600 | 0.222 | -0.586 | 0.00127 | 3.18447 | 5 | 74.5 | 0.15 | 5 | null | 1 | 50 | 10 | ; | 10 | 20 | ; | 20 | null |
| Atlantic bonito | *Sarda sarda* | 82.380 | 0.517 | -1.548 | 0.00656 | 3.11781 | 65 | 40.0 | 0.27 | 4 | 24.5 | null | 29435 | 5 | ; | 5 | 30 | ; | 9 | 10 |
| European pilchard | *Sardina pilchardus* | 21.302 | 0.445 | -0.852 | 0.00638 | 3.08475 | 350 | 13.4 | 0.45 | 5.5 | 11 | null | 212808 | 15 | ; | 15 | 500 | ; | 700 | 10 |
| Round sardinella | *Sardinella aurita* | 27.538 | 0.437 | -2.100 | 0.01112 | 2.96122 | 415 | 15.2 | 0.10 | 6.33 | 10 | null | 24628 | 10 | ; | 10 | 500 | ; | 700 | 10 |
| Brushtooth lizardfish | *Saurida undosquamis* | 41.765 | 0.232 | -0.589 | 0.00740 | 2.97890 | 264 | 16.3 | 0.50 | 7.5 | null | 1 | 74 | 6 | ; | 6 | 85 | ; | 85 | null |
| Brown meagre | *Sciaena umbra* | 54.050 | 0.184 | -0.394 | 0.02140 | 3.05940 | 279 | 25.8 | 0.21 | 21 | null | 1 | 697 | 6 | ; | 6 | 95 | ; | 95 | null |
| Chub mackerel | *Scomber japonicus* | 39.925 | 0.315 | -0.854 | 0.00615 | 3.22250 | 1567 | 29.2 | 0.19 | 10.5 | 15 | null | 12902 | 6 | ; | 6 | 50 | ; | 50 | null |
| Atlantic mackerel | *Scomber scombrus* | 37.300 | 0.431 | -0.925 | 0.00961 | 3.07767 | 415 | 20.8 | 0.19 | 17 | 15 | null | 9081 | 5 | ; | 5 | 50 | ; | 50 | null |
| Turbot | *Scophthalmus maximus* | 63.064 | 0.290 | -0.408 | 0.02869 | 2.99830 | 1078 | 46.3 | 0.25 | 5.3 | null | 1 | 1466 | 19.3 | ; | 19.3 | 172.7 | ; | 172.7 | null |
| Small red scorpionfish | *Scorpaena notata* | 16.925 | 0.310 | -1.810 | 0.01776 | 3.02234 | 197 | 11.5 | 0.20 | 8 | null | 1 | 0 | 4.5 | ; | 4.5 | 20 | ; | 20 | null |
| Lesser spotted dogfish | *Scyliorhinus canicula* | 56.800 | 0.530 | 0.000 | 0.00153 | 3.19565 | 182 | 41.3 | 0.20 | 10.5 | 25 | null | 440 | 5 | ; | 5 | 50 | ; | 50 | null |
| Common cuttlefish | *Sepia officinalis* | 28.383 | 0.800 | -0.060 | 0.29750 | 2.64750 | 7 | 11.0 | 0.20 | 1.5 | 5 | null | 14554 | 5 | ; | 5 | 30 | ; | 30 | null |
| Greater amberjack | *Seriola dumerili* | 174.600 | 0.190 | -0.314 | 0.02120 | 2.90550 | 173 | 104 | 0.31 | 15 | null | 1 | 2975 | 4.5 | ; | 4.5 | 12 | ; | 12 | null |
| Blacktail comber | *Serranus atricauda* | 49.500 | 0.110 | -0.760 | 0.00905 | 3.17550 | 5655 | 19.3 | 0.31 | 16 | null | 1 | 0 | 6 | ; | 6 | 124 | ; | 124 | null |
| Common sole | *Solea solea* | 39.600 | 0.440 | -0.460 | 0.00700 | 3.06380 | 500 | 20.0 | 0.20 | 16.7 | 16 | null | 6020 | 22 | ; | 22 | 330 | ; | 330 | null |
| Gilthead seabream | *Sparus aurata* | 57.033 | 0.284 | -0.627 | 0.02598 | 3.01966 | 108 | 32.2 | 0.05 | 11 | 15 | null | 12004 | 6 | ; | 6 | 45 | ; | 45 | null |
| European barracuda | *Sphyraena sphyraena* | 55.300 | 0.123 | -3.248 | 0.05686 | 2.52700 | 70 | 24.5 | 0.22 | 8 | null | 1 | 2245 | 3 | ; | 3 | 8 | ; | 8 | null |
| Yellowmouth barracuda | *Sphyraena viridensis* | 100.600 | 0.089 | -0.825 | 0.00391 | 3.00000 | 56 | 61.0 | 0.22 | 8 | null | 1 | 0 | 3 | ; | 3 | 15 | ; | 15 | null |
| Blotched picarel | *Spicara maena* | 21.990 | 0.255 | -1.160 | 0.00280 | 3.50500 | 350 | 11.7 | 0.10 | 5 | 13 | null | 1402 | 8 | ; | 8 | 20 | ; | 20 | null |
| Picarel | *Spicara smaris* | 19.600 | 0.230 | -1.970 | 0.01134 | 2.93105 | 349 | 9.1 | 0.20 | 5.5 | 9 | null | 5273 | 11 | ; | 11 | 30 | ; | 30 | null |
| Black seabream | *Spondyliosoma cantharus* | 41.700 | 0.161 | -0.848 | 0.03121 | 2.94075 | 664 | 22.0 | 0.42 | 13 | null | 1 | 1101 | 6 | ; | 6 | 45 | ; | 45 | null |

| | | | | | | | | | | | | | | | | | | | | |
|---|---|---|---|---|---|---|---|---|---|---|---|---|---|---|---|---|---|---|---|---|
| European sprat | *Sprattus sprattus* | 14.296 | 0.344 | -1.399 | 0.00866 | 2.94760 | 246 | 10.8 | 0.60 | 6 | 7.5 | null | 992 | 8 | ; | 12 | 100 | ; | 100 | null |
| Spottail mantis shrimp | *Squilla mantis* | 19.690 | 0.500 | -0.370 | 0.09900 | 1.73700 | 1618 | 14.0 | 0.50 | 3.5 | 1.25 | null | 7926 | 5 | ; | 9 | 100 | ; | 50 | 2 |
| Reticulated leatherjacket | *Stephanolepis diaspros* | 27.830 | 0.350 | -0.499 | 0.01753 | 3.03367 | 2589 | 9.3 | 0.20 | 4 | null | 1 | 0 | 15 | ; | 15 | 30 | ; | 30 | null |
| Albacore | *Thunnus alalunga* | 94.700 | 0.258 | -1.354 | 0.00339 | 2.88000 | 130 | 66.3 | 0.85 | 6 | 39 | null | 3896 | 2.78 | ; | 2.78 | 41.04 | ; | 41.04 | null |
| Atlantic bluefin tuna | *Thunnus thynnus* | 319.000 | 0.093 | -0.970 | 0.02310 | 2.93400 | 93 | 111 | 0.37 | 14 | 50 | null | 16405 | 2.58 | ; | 2.58 | 21.46 | ; | 21.46 | null |
| Mediterranean horse mackerel | *Trachurus mediterraneus* | 35.209 | 0.335 | -0.773 | 0.01140 | 2.97076 | 900 | 18.0 | 0.01 | 12 | 11.5 | null | 51710 | 5 | ; | 5 | 40 | ; | 40 | null |
| Blue jack mackerel | *Trachurus picturatus* | 62.700 | 0.080 | -2.820 | 0.00890 | 2.96000 | 1018 | 24.8 | 0.20 | 18 | 11 | null | 787 | 6 | ; | 6 | 40 | ; | 40 | null |
| Atlantic horse mackerel | *Trachurus trachurus* | 32.265 | 0.322 | -0.709 | 0.00956 | 3.00361 | 893 | 16.5 | 0.05 | 9 | 9.25 | null | 48019 | 5 | ; | 5 | 40 | ; | 40 | null |
| Piper gurnard | *Trigla lyra* | 61.466 | 0.162 | -1.028 | 0.01197 | 2.91925 | 178 | 19.0 | 0.20 | 7.5 | null | 1 | 24 | 6 | ; | 6 | 85 | ; | 85 | null |
| Pouting | *Trisopterus luscus* | 44.375 | 0.351 | -0.615 | 0.01086 | 3.12462 | 473 | 21.5 | 0.05 | 5 | null | 1 | 742 | 5 | ; | 5 | 9.67 | ; | 9.67 | null |
| Poor cod | *Trisopterus minutus* | 25.943 | 0.399 | -2.144 | 0.00754 | 3.14413 | 76 | 13.3 | 0.10 | 4.33 | 11 | null | 2377 | 2.2 | ; | 2.2 | 10.67 | ; | 10.67 | null |
| Goldband goatfish | *Upeneus moluccensis* | 25.698 | 0.830 | 0.000 | 0.01027 | 3.21717 | 6433 | 11.0 | 0.20 | 5 | null | 1 | 61 | 5 | ; | 5 | 25 | ; | 25 | null |
| Swordfish | *Xiphias gladius* | 238.500 | 0.185 | -1.404 | 0.00475 | 3.17100 | 40 | 223 | 0.20 | 10 | 50 | null | 14377 | 2 | ; | 2 | 24.18 | ; | 24.18 | null |
| Grass goby | *Zosterisessor ophiocephalus* | 27.400 | 0.169 | -2.120 | 0.00870 | 3.10233 | 500 | 11.8 | 0.20 | 5 | null | 1 | 0 | 9 | ; | 9 | 35 | ; | 35 | null |
| Euphausiids | - | 1.840 | 1.680 | -0.198 | 0.00900 | 2.92000 | 42254 | 1.0 | 0.25 | 2 | null | 1 | 0 | 100 | ; | 100 | 400 | ; | 200 | 0.6 |

#### Appendix C. Reference list of input parameters.

Table C.1. Reference list of input parameters

| Common name | Scientific name | Growth | Size/Age at maturity | Reproduction | Spawning period | Egg size | Feeding | Catches | Biomass | Source of growth data |
|---|---|---|---|---|---|---|---|---|---|---|
| Allis shad | *Alosa alosa* | Froese and Pauly, 2017 | Froese and Pauly, 2017 | Aprahamian et al, 2003 | Aprahamian et al, 2003 | By default | Barnes et al, 2008 | Mean FAO (GFCM) and Sea Around Us | | Other |
| Twaite shad | *Alosa fallax* | Froese and Pauly, 2017 | Froese and Pauly, 2017 | Aprahamian et al, 2003 | Aprahamian et al, 2003 | By default | Barnes et al, 2008 | Mean FAO (GFCM) and Sea Around Us | | Med Sea |
| European eel | *Anguilla anguilla* | Froese and Pauly, 2017 | Froese and Pauly, 2017 | MacNamara et al, 2016 | MacNamara et al, 2016 | By default | Barnes et al, 2008 | Mean FAO (GFCM) and Sea Around Us | | Med Sea |
| Meagre | *Argyrosomus regius* | Froese and Pauly, 2017 | Froese and Pauly, 2018 | Gil et al, 2013 | Gil et al, 2013 | By default | Barnes et al, 2008 | Mean FAO (GFCM) and Sea Around Us | | Other |
| Giant red shrimp | *Aristaeomorpha foliacea* | STECF 2015b | STECF 2015b | Kapiris and Thessalou-Legaki, 2006 | Kapiris and Thessalou-Legaki, 2006 | By default | Barnes et al, 2008 | Mean FAO (GFCM) and Sea Around Us | http://www.fao.org/gfcm/data/safs | Med Sea |
| Blue and red shrimp | *Aristeus antennatus* | STECF 2015b | STECF 2015b | Kapiris and Thessalou-Legaki, 2006 | Kapiris and Thessalou-Legaki, 2006 | By default | Barnes et al, 2008 | Mean FAO (GFCM) and Sea Around Us | | Med Sea |
| Big-scale sand smelt | *Atherina boyeri* | Froese and Pauly, 2017 | Tsikliras and Stergiou, 2013 | Patimar et al, 2009 | Tsikliras et al, 2010 | By default | Barnes et al, 2008 | Mean FAO (GFCM) and Sea Around Us | | Med Sea |
| Bullet tuna | *Auxis rochei rochei* | Kahraman et al, 2011 | Kahraman et al, 2011 | Macias et al, 2006 | Macias et al, 2006 | Neuheimer et al, 2016 | Barnes et al, 2008 | Mean FAO (GFCM) and Sea Around Us | | Med Sea |
| Garfish | *Belone belone* | Froese and Pauly, 2017 | Tsikliras and Stergiou, 2013 | Zorica et al, 2010 | Tsikliras et al, 2010 | By default | Barnes et al, 2008 | Mean FAO (GFCM) and Sea Around Us | | Med Sea |
| Bogue | *Boops boops* | STECF 2012 | STECF 2012 | Lamrini, 2010 | Tsikliras et al, 2010 | By default | Barnes et al, 2008 | Mean FAO (GFCM) and Sea Around Us | | Med Sea |
| Blue runner | *Caranx crysos* | Froese and Pauly, 2017 | Tsikliras and Stergiou, 2013 | Froese and Pauly, 2017 | Tsikliras et al, 2010 | By default | Barnes et al, 2008 | Mean FAO (GFCM) and Sea Around Us | | Other |
| Tub gurnard | *Chelidonichthys lucerna* | Froese and Pauly, 2017 | Tsikliras and Stergiou, 2013 | Cicek et al, 2008 | Tsikliras et al, 2010 | By default | Barnes et al, 2008 | Mean FAO (GFCM) and Sea Around Us | | Med Sea |
| Mediterranean rainbow wrasse | *Coris julis* | Froese and Pauly, 2017 | Froese and Pauly, 2018 | Alonso-Fernandez et al, 2013 | Tsikliras et al, 2010 | Alonso-Fernandez et al, 2013 | Barnes et al, 2008 | Mean FAO (GFCM) and Sea Around Us | | Med Sea |
| Common dolphin fish | *Coryphaena hippurus* | STECF 2014 | STECF 2014 | Massuti and Morales-Nin, 1997 | Tsikliras et al, 2010 | By default | Barnes et al, 2008 | Mean FAO (GFCM) and Sea Around Us | | Med Sea |
| Common shrimp | *Crangon crangon* | Froese and Pauly, 2017 Kasapoglu et Duzgunes, 2013 | Bilgin and Samsun, 2006 | Bilgin and Samsun, 2006 | Bilgin and Samsun, 2006 | Bilgin and Samsun, 2006 | Barnes et al, 2008 | Mean FAO (GFCM) and Sea Around Us | | Other |
| Cristal goby | *Crystallogobius linearis* | Froese and Pauly, 2017 | Tsikliras and Stergiou, 2013 | Caputo et al, 2003 | Tsikliras et al, 2010 | Caputo et al, 2003 | Barnes et al, 2008 | Mean FAO (GFCM) and Sea Around Us | | From other ecosytem |
| Common dentex | *Dentex dentex* | Froese and Pauly, 2017 | Tsikliras and Stergiou, 2013 | Loir et al, 2001 | Tsikliras et al, 2010 | By default | Barnes et al, 2008 | Mean FAO (GFCM) and Sea Around Us | | Med Sea |
| Pink dentex | *Dentex gibbosus* | Froese and Pauly, 2017 | Tsikliras and Stergiou, 2013 | Grubisic et al, 2007 | Tsikliras et al, 2010 | By default | Barnes et al, 2008 | Mean FAO (GFCM) and Sea Around Us | | Med Sea |
| Morocco dentex | *Dentex maroccanus* | Froese and Pauly, 2017 | Froese and Pauly, 2018 | Lamrini and Bouymajjane, 2011 | Lamrini and Bouymajjane, 2011 | By default | Barnes et al, 2008 | Mean FAO (GFCM) and Sea Around Us | | Med Sea |

| | | | | | | | | | | |
|---|---|---|---|---|---|---|---|---|---|---|
| European seabass | *Dicentrarchus labrax* | Froese and Pauly, 2017 | Tsikliras and Stergiou, 2013 | Mayer et al, 1990<br>Wassef and Emary, 1989<br>Kara, 1997 | Tsikliras et al, 2010 | Neuheimer et al, 2016 | Barnes et al, 2008 | Mean FAO (GFCM) and Sea Around Us | | Med Sea |
| Annular sea bream | *Diplodus annularis* | Froese and Pauly, 2017 | Tsikliras and Stergiou, 2013 | Chaouch et al, 2013 | Tsikliras et al, 2010 | Chaouch et al, 2013 | Barnes et al, 2008 | Mean FAO (GFCM) and Sea Around Us | | Med Sea |
| Zebra seabream | *Diplodus cervinus* | Froese and Pauly, 2017 | Tsikliras and Stergiou, 2013 | Derbal, 2010 | Derbal, 2010 | By default | Barnes et al, 2008 | Mean FAO (GFCM) and Sea Around Us | | From other ecosytem |
| Sharpsnout seabream | *Diplodus puntazzo* | Froese and Pauly, 2017 | Tsikliras and Stergiou, 2013 | Papadaki et al, 2008 | Tsikliras et al, 2010 | By default | Barnes et al, 2008 | Mean FAO (GFCM) and Sea Around Us | | Med Sea |
| White sea bream | *Diplodus sargus sargus* | Froese and Pauly, 2017 | Tsikliras and Stergiou, 2013 | Mouine et al, 2007 | Tsikliras et al, 2010 | By default | Barnes et al, 2008 | Mean FAO (GFCM) and Sea Around Us | | Med Sea |
| Common two-banded sea bream | *Diplodus vulgaris* | Froese and Pauly, 2017 | Tsikliras and Stergiou, 2013 | Hadj Taieb et al, 2013 | Tsikliras et al, 2010 | By default | Barnes et al, 2008 | Mean FAO (GFCM) and Sea Around Us | | Med Sea |
| Horned octopus | *Eledone cirrhosa* | Froese and Pauly, 2017 | Rjeibi et al, 2014 | Rjeibi et al, 2014 | Rjeibi et al, 2014 | By default | Barnes et al, 2008 | Mean FAO (GFCM) and Sea Around Us | | Med Sea |
| European anchovy | *Engraulis encrasicolus* | STECF 2015b | STECF 2015b | Somarakis et al, 2004 | Tsikliras et al, 2010 | Somarakis et al, 2004 | Barnes et al, 2008 | Mean FAO (GFCM) and Sea Around Us | http://www.fao.org/gfcm/data/safs | Med Sea |
| White grouper | *Epinephelus aeneus* | Froese and Pauly, 2017 | Froese and Pauly, 2018 | Bouain and Siau, 1983 | Bouain and Siau, 1983 | By default | Barnes et al, 2008 | Mean FAO (GFCM) and Sea Around Us | | Med Sea |
| Dusky grouper | *Epinephelus marginatus* | Froese and Pauly, 2017 | Tsikliras and Stergiou, 2013 | Renones et al, 2010 | Tsikliras et al, 2010 | By default | Barnes et al, 2008 | Mean FAO (GFCM) and Sea Around Us | | Med Sea |
| Red-eye round herring | *Etrumeus teres* | Yilmaz and Hosucu, 2003 | Osman et al, 2011 | Osman et al, 2011 | Tsikliras et al, 2010 | Osman et al, 2011 | Barnes et al, 2008 | Mean FAO (GFCM) and Sea Around Us | | Med Sea |
| Grey gurnard | *Eutrigla gurnardus* | Froese and Pauly, 2017 | Tsikliras and Stergiou, 2013 | Gokçe, 1998 | Tsikliras et al, 2010 | By default | Barnes et al, 2008 | Mean FAO (GFCM) and Sea Around Us | | Med Sea |
| Black-mouthed dogfish | *Galeus melastomus* | STECF 2011 | STECF 2011 | Capapé et al, 2008 | Tsikliras et al, 2010 | Capapé et al, 2008 | Barnes et al, 2008 | Mean FAO (GFCM) and Sea Around Us | | Med Sea |
| Black goby | *Gobius niger* | Froese and Pauly, 2017 | Bouchereau, 1997 | Bouchereau, 1997 | Tsikliras et al, 2010 | By default | Barnes et al, 2008 | Mean FAO (GFCM) and Sea Around Us | | Med Sea |
| Lusitanian toadfish | *Halobatrachus didactylus* | Froese and Pauly, 2017 | Palazon-Fernandez et al, 2001 | Palazon-Fernandez et al, 2001 | Palazon-Fernandez et al, 2001 | By default | Barnes et al, 2008 | Mean FAO (GFCM) and Sea Around Us | | From other ecosytem |
| Shortfin squid | *Illex coindetii* | Palomares and Pauly, 2017<br>Belcari, 1996<br>Sanchez, 1984 | Gonzalez et al, 1996 | Gonzalez et al, 1996<br>Laptikhovsky et Nigmatullin et al, 1993 | Gonzalez et al, 1996 | Neuheimer et al, 2016 | Barnes et al, 2008 | Mean FAO (GFCM) and Sea Around Us | | Other |
| Megrim | *Lepidorhombus whiffiagonis* | Froese and Pauly, 2017 | Tsikliras and Stergiou, 2013 | Macdonald, 2014 | Tsikliras et al, 2010 | Neuheimer et al, 2016 | Barnes et al, 2008 | Mean FAO (GFCM) and Sea Around Us | | From other ecosytem |
| Golden grey mullet | *Liza aurata* | Froese and Pauly, 2017 | Tsikliras and Stergiou, 2013 | Hotos at al, 2000<br>Abdallah et al, 2013 | Tsikliras et al, 2010 | By default | Barnes et al, 2008 | Mean FAO (GFCM) and Sea Around Us | | Med Sea |
| Thinlip grey mullet | *Liza ramada* | Froese and Pauly, 2017 | Tsikliras and Stergiou, 2013 | Fazli et al, 2008 | Tsikliras et al, 2010 | By default | Barnes et al, 2008 | Mean FAO (GFCM) and Sea Around Us | | Med Sea |
| Leaping mullet | *Liza saliens* | Froese and Pauly, 2017 | Tsikliras and Stergiou, 2013 | Koutrakis, 2011 | Tsikliras et al, 2010 | By default | Barnes et al, 2008 | Mean FAO (GFCM) and Sea Around Us | | Med Sea |
| European squid | *Loligo vulgaris* | Froese and Pauly, 2017 | Laptikhovsky, 2000 | Laptikhovsky, 2000 | Laptikhovsky, 2000 | Laptikhovsky, 2000 | Barnes et al, 2008 | Mean FAO (GFCM) and Sea Around Us | | Med Sea |
| Black-bellied angler | *Lophius budegassa* | STECF 2015a | STECF 2015a | Colmenero et al, 2013 | Tsikliras et al, 2010 | Colmenero et al, 2013 | Barnes et al, 2008<br>Scharf et al, 2000 | Mean FAO (GFCM) and Sea Around Us | http://www.fao.org/gfcm/data/safs | Med Sea |

| | | | | | | | | | | |
|---|---|---|---|---|---|---|---|---|---|---|
| Anglerfish | *Lophius piscatorius* | STECF 2015a | STECF 2015a | Colmenero et al, 2013 | Tsikliras et al, 2010 | Colmenero et al, 2013 | Barnes et al, 2008 Scharf et al, 2000 | Mean FAO (GFCM) and Sea Around Us | | Med Sea |
| Whiting | *Merlangius merlangus* | Froese and Pauly, 2017 | Tsikliras and Stergiou, 2013 | Hislop and Hall, 1974 | Tsikliras et al, 2010 | Neuheimer et al, 2016 | Barnes et al, 2008 | Mean FAO (GFCM) and Sea Around Us | | Med Sea |
| European hake | *Merluccius merluccius* | STECF 2015b | STECF 2015b | Recasens et al, 2008 Al-Absawy et al, 2010 | Tsikliras et al, 2010 | Recasens et al, 2008 | Barnes et al, 2008 | Mean FAO (GFCM) and Sea Around Us | http://www.fao.org/gfcm/data/safs | Med Sea |
| Blue whiting | *Micromesistius poutassou* | STECF 2014 | STECF 2014 | Macchi et al, 2005 | Tsikliras et al, 2010 | By default | Barnes et al, 2008 | Mean FAO (GFCM) and Sea Around Us | | Med Sea |
| Flathead grey mullet | *Mugil cephalus* | Froese and Pauly, 2017 | Tsikliras and Stergiou, 2013 | Froese and Pauly, 2017 | Tsikliras et al, 2010 | By default | Barnes et al, 2008 Scharf et al, 2000 | Mean FAO (GFCM) and Sea Around Us | | Med Sea |
| Red mullet | *Mullus barbatus barbatus* | STECF 2015a | STECF 2015a | Tirasin et al, 2007 Layachi et al, 2007 | Tsikliras et al, 2010 | Layachi et al, 2007 | Barnes et al, 2008 | Mean FAO (GFCM) and Sea Around Us | http://www.fao.org/gfcm/data/safs | Med Sea |
| Striped red mullet | *Mullus surmuletus* | STECF 2013 | STECF 2013 | Amin et al, 2016 | Tsikliras et al, 2010 | Amin et al, 2016 | Barnes et al, 2008 | Mean FAO (GFCM) and Sea Around Us | | Med Sea |
| Smooth hound | *Mustelus mustelus* | Froese and Pauly, 2017 | Tsikliras and Stergiou, 2013 | Saidi, 2008 | Saidi, 2008 | Saidi, 2008 | Barnes et al, 2008 Scharf et al, 2000 | Mean FAO (GFCM) and Sea Around Us | | Med Sea |
| Norway lobster | *Nephrops norvegicus* | STECF 2015a | STECF 2015a | Briggs et al, 2002 | Briggs et al, 2002 | By default | Barnes et al, 2008 | Mean FAO (GFCM) and Sea Around Us | | Med Sea |
| Common octopus | *Octopus vulgaris* | Froese and Pauly, 2017 | Zghidi et al, 2004 Cuccu et al, 2013 | Zghidi et al, 2004 Cuccu et al, 2013 | Zghidi et al, 2004 Cuccu et al, 2013 | Cuccu et al, 2013 | Barnes et al, 2008 | Mean FAO (GFCM) and Sea Around Us | | Med Sea |
| Axillary seabream | *Pagellus acarne* | Froese and Pauly, 2017 | Tsikliras and Stergiou, 2013 | Velasco et al, 2011 | Tsikliras et al, 2010 | By default | Barnes et al, 2008 | Mean FAO (GFCM) and Sea Around Us | | Med Sea |
| Common pandora | *Pagellus erythrinus* | Froese and Pauly, 2017 | Tsikliras and Stergiou, 2013 | Klaoudatos et al, 2004 | Tsikliras et al, 2010 | By default | Barnes et al, 2008 | Mean FAO (GFCM) and Sea Around Us | | Med Sea |
| Common seabream | *Pagrus pagrus* | Pajuelo and Lorenzo, 1996 | Mylonas et al, 2004 Aristizabal, 2009 | Mylonas et al, 2004 Aristizabal, 2009 | Mylonas et al, 2004 Aristizabal, 2009 | Aristizabal, 2009 | Barnes et al, 2008 | Mean FAO (GFCM) and Sea Around Us | | Med Sea |
| Common prawn | *Palaemon serratus* | Froese and Pauly, 2017 | Bilgin and Samsun, 2006 | Bilgin and Samsun, 2006 | Bilgin and Samsun, 2006 | By default | Barnes et al, 2008 | Mean FAO (GFCM) and Sea Around Us | | Other |
| Common spiny lobster | *Palinurus elephas* | Froese and Pauly, 2017 | Goni et al, 2003 | Goni et al, 2003 | Goni et al, 2003 | Goni et al, 2003 | Barnes et al, 2008 | Mean FAO (GFCM) and Sea Around Us | | Other |
| Deep-water rose shrimp | *Parapenaeus longirostris* | STECF 2015a | STECF 2015a | Sobrino and Garcia, 2007 | Sobrino and Garcia, 2007 | By default | Barnes et al, 2008 | Mean FAO (GFCM) and Sea Around Us | http://www.fao.org/gfcm/data/safs | Med Sea |
| Caramote prawn | *Penaeus kerathurus* | Ben Meriem et al, 2004 | Ben Meriem et al, 2004 | Jawadi and Ben Mariem, 2007 | Jawadi and Ben Mariem, 2007 | Lumare et al, 2011 | Barnes et al, 2008 | Mean FAO (GFCM) and Sea Around Us | | Med Sea |
| Forkbeard | *Phycis phycis* | Froese and Pauly, 2017 | Vieira et al, 2016 | Vieira et al, 2016 | Vieira et al, 2016 | By default | Barnes et al, 2008 | Mean FAO (GFCM) and Sea Around Us | | Med Sea |
| European flounder | *Platichthys flesus flesus* | Froese and Pauly, 2017 | Aydin et al, 2011 | Aydin et al, 2011 | Tsikliras et al, 2010 | Aydin et al, 2011 | Barnes et al, 2008 | Mean FAO (GFCM) and Sea Around Us | | Med Sea |
| European plaice | *Pleuronectes platessa* | Froese and Pauly, 2017 | Froese and Pauly, 2017 | Froese and Pauly, 2017 | Froese and Pauly, 2017 | Froese and Pauly, 2017 | Barnes et al, 2008 | Mean FAO (GFCM) and Sea Around Us | | Other |

| | | | | | | | | | | |
|---|---|---|---|---|---|---|---|---|---|---|
| Bluefish | *Pomatomus saltatrix* | Froese and Pauly, 2017 | Tsikliras and Stergiou, 2013 | Villegas-Hernandez, 2015 | Tsikliras et al, 2010 | By default | Barnes et al, 2008 Scharf et al, 2000 | Mean FAO (GFCM) and Sea Around Us | | Other |
| Marbled goby | *Pomatoschistus marmoratus* | Froese and Binohan 2000 | Tsikliras and Stergiou, 2013 | Mazzoldi et al, 2002 | Tsikliras et al, 2010 | Mazzoldi et al, 2002 | Barnes et al, 2008 | Mean FAO (GFCM) and Sea Around Us | | Other |
| Sand goby | *Pomatoschistus minutus* | Froese and Pauly, 2017 | Bouchereau, 1997 | Bouchereau, 1997 | Tsikliras et al, 2010 | By default | Barnes et al, 2008 | Mean FAO (GFCM) and Sea Around Us | | Other |
| Common guitarfish | *Rhinobatos rhinobatos* | Froese and Pauly, 2017 | Tsikliras and Stergiou, 2013 | Enajjar, 2008 | Tsikliras et al, 2010 | Enajjar, 2008 | Barnes et al, 2008 | Mean FAO (GFCM) and Sea Around Us | | Med Sea |
| Atlantic bonito | *Sarda sarda* | Froese and Pauly, 2017 | | Macias et al, 2005 | Tsikliras et al, 2010 | By default | Barnes et al, 2008 | ICCAT 2016 | | Med Sea |
| European pilchard | *Sardina pilchardus* | STECF 2015a | STECF 2015 | Somarakis et al, 2002 | Tsikliras et al, 2010 | Somarakis et al, 2002 | Barnes et al, 2008 | Mean FAO (GFCM) and Sea Around Us | http://www.fao.org/gfcm/data/safs | Med Sea |
| Round sardinella | *Sardinella aurita* | Froese and Pauly, 2017 | Tsikliras and Stergiou, 2013 | Tsikliras and Antonopoulou, 2006 Mustac and Sinovcic, 2012 | Tsikliras et al, 2010 | Mustac and Sinovcic, 2012 | Barnes et al, 2008 | Mean FAO (GFCM) and Sea Around Us | | Med Sea |
| Brushtooth lizardfish | *Saurida undosquamis* | GFCM 2014 | GFCM 2014 | Kadharsha et al, 2013 | Tsikliras et al, 2010 | By default | Barnes et al, 2008 | Mean FAO (GFCM) and Sea Around Us | | Med Sea |
| Brown meagre | *Sciaena umbra* | Froese and Pauly, 2017 | Tsikliras and Stergiou, 2013 | Derbal and Kara, 2007 | Tsikliras et al, 2010 | By default | Barnes et al, 2008 | Mean FAO (GFCM) and Sea Around Us | | Med Sea |
| Chub mackerel | *Scomber japonicus* | Froese and Pauly, 2017 | Techetach et al, 2010 | Techetach et al, 2010 | Tsikliras et al, 2010 | Techetach et al, 2010 | Barnes et al, 2008 | Mean FAO (GFCM) and Sea Around Us | | Med Sea |
| Atlantic mackerel | *Scomber scombrus* | Froese and Pauly, 2017 | Tsikliras and Stergiou, 2013 | Meneghesso et al, 2013 | Tsikliras et al, 2010 | Neuheimer et al, 2016 | Barnes et al, 2008 | Mean FAO (GFCM) and Sea Around Us | | Med Sea |
| Turbot | *Scophthalmus maximus* | Froese and Pauly, 2017 | Tsikliras and Stergiou, 2013 | Meneghesso et al, 2013 | Meneghesso et al, 2013 | By default | Barnes et al, 2008 Scharf et al, 2000 | Mean FAO (GFCM) and Sea Around Us | | Med Sea |
| Small red scorpionfish | *Scorpaena notata* | Froese and Pauly, 2017 | Tsikliras and Stergiou, 2013 | Munoz et al, 2005 | Tsikliras et al, 2010 | By default | Barnes et al, 2008 | Mean FAO (GFCM) and Sea Around Us | | Med Sea |
| Lesser spotted dogfish | *Scyliorhinus canicula* | Froese and Pauly, 2017 | Tsikliras and Stergiou, 2013 | Capapé et al, 2008 | Capapé et al, 2008 | Capapé et al, 2008 | Barnes et al, 2008 | Mean FAO (GFCM) and Sea Around Us | | Med Sea |
| Common cuttlefish | *Sepia officinalis* | Palomares and Pauly, 2017 | Ezzeddine-Najai, 1993 | Ezzeddine-Najai, 1993; Guera, 2006 | Ezzeddine-Najai, 1993; Guera, 2006 | Ezzeddine-Najai, 1993; Guera, 2006 | Barnes et al, 2008 | Mean FAO (GFCM) and Sea Around Us | | Med Sea |
| Greater amberjack | *Seriola dumerili* | Froese and Pauly, 2017 | Honebrink, 2000 | Honebrink, 2000 | Honebrink, 2000 | By default | Barnes et al, 2008 | Mean FAO (GFCM) and Sea Around Us | | Med Sea |
| Blacktail comber | *Serranus atricauda* | Froese and Pauly, 2017 | Tsikliras and Stergiou, 2013 | Neves et al, 2013 | Neves et al, 2013 | Neves et al, 2013 | Barnes et al, 2008 | Mean FAO (GFCM) and Sea Around Us | | Other |
| Common sole | *Solea solea* | STECF 2011 | STECF 2011 | Farrugio and Le Corre, 1986 | Tsikliras et al, 2010 | Neuheimer et al, 2016 | Barnes et al, 2008 | Mean FAO (GFCM) and Sea Around Us | | Med Sea |
| Gilthead seabream | *Sparus aurata* | Froese and Pauly, 2017 | Meneghesso et al, 2013 | Meneghesso et al, 2013 | Tsikliras et al, 2010 | Jeres et al, 2012 | Barnes et al, 2008 | Mean FAO (GFCM) and Sea Around Us | | Med Sea |
| European barracuda | *Sphyraena sphyraena* | Froese and Pauly, 2017 | Tsikliras and Stergiou, 2013 | Meneghesso et al, 2013 | Tsikliras et al, 2010 | By default | Barnes et al, 2008 | Mean FAO (GFCM) and Sea Around Us | | Med Sea |
| Yellowmouth barracuda | *Sphyraena viridensis* | Froese and Pauly, 2017 | Tsikliras and Stergiou, 2013 | Meneghesso et al, 2013 | Meneghesso et al, 2013 | By default | Barnes et al, 2008 | Mean FAO (GFCM) and Sea Around Us | | Med Sea |
| Blotched picarel | *Spicara maena* | Froese and Pauly, 2017 | Tsikliras and Stergiou, 2013 | Matic-Skoko et al, 2004 | Tsikliras et al, 2010 | By default | Barnes et al, 2008 | Mean FAO (GFCM) and Sea Around Us | | Med Sea |
| Picarel | *Spicara smaris* | STECF 2012 | STECF 2012 | karlou-Riga et al, 2007 | Tsikliras et al, 2010 | By default | Barnes et al, 2008 | Mean FAO (GFCM) and Sea Around Us | | Med Sea |
| Black seabream | *Spondyliosoma cantharus* | Froese and Pauly, 2017 | Tsikliras and Stergiou, 2013 | Dulcic et al, 1998 | Tsikliras et al, 2010 | Neuheimer et al, 2016 | Barnes et al, 2008 | Mean FAO (GFCM) and Sea Around Us | | Med Sea |

| | | | | | | | | | | |
|---|---|---|---|---|---|---|---|---|---|---|
| European sprat | *Sprattus sprattus* | Froese and Pauly, 2017 | Alheit, 1988 | Alheit, 1988 | Tsikliras et al, 2010 | Neuheimer et al, 2016 | Barnes et al, 2008 | Mean FAO (GFCM) and Sea Around Us | | Med Sea |
| Spottail mantis shrimp | *Squilla mantis* | STECF 2011 | STECF 2012 | Mili et al, 2014 | Mili et al, 2014 | By default | Barnes et al, 2008 | Mean FAO (GFCM) and Sea Around Us | | Med Sea |
| Reticulated leatherjacket | *Stephanolepis diaspros* | Box, 2008 | Tsikliras and Stergiou, 2013 | Rim and Mohamed-Nejmeddine, 2011 | Rim and Mohamed-Nejmeddine, 2011 | By default | Barnes et al, 2008 | Mean FAO (GFCM) and Sea Around Us | | Med Sea |
| Albacore | *Thunnus alalunga* | ICCAT 2017 | ICCAT 2017 | Saber et al, 2011 | Saber et al, 2011 | By default | Barnes et al, 2008 Ménard et al, 2006 | ICCAT 2016 | Winker Henning (comm. pers.) | Med Sea |
| Atlantic bluefin tuna | *Thunnus thynnus* | ICCAT 2017 | ICCAT 2017 | Medina et al, 2007 Fromentin and Powers, 2005 | ICCAT 2016 | By default | Barnes et al, 2008 Ménard et al, 2006 | ICCAT 2016 | Fromentin Jean-Marc (comm. pers.) | Med Sea |
| Mediterranean horse mackerel | *Trachurus mediterraneus* | Froese and Pauly, 2017 | Tsikliras and Stergiou, 2013 | Demirel and Yuksek, 2013 | Tsikliras et al, 2010 | By default | Barnes et al, 2008 | Mean FAO (GFCM) and Sea Around Us | | Med Sea |
| Blue jack mackerel | *Trachurus picturatus* | Froese and Pauly, 2017 | Tsikliras and Stergiou, 2013 | Vasconcelos et al, 2017 | Vasconcelos et al, 2017 | By default | Barnes et al, 2008 | Mean FAO (GFCM) and Sea Around Us | | Med Sea |
| Atlantic horse mackerel | *Trachurus trachurus* | Froese and Pauly, 2017 | Tsikliras and Stergiou, 2013 | Van Damme et al, 2013 Gherram et al, 2013 Abaunza et al, 2003 | Tsikliras et al, 2010 | By default | Barnes et al, 2008 | Mean FAO (GFCM) and Sea Around Us | | Med Sea |
| Piper gurnard | *Trigla lyra* | Froese and Pauly, 2017 | Tsikliras and Stergiou, 2013 | Agbali et al, 2015 | Tsikliras et al, 2010 | By default | Barnes et al, 2008 | Mean FAO (GFCM) and Sea Around Us | | Med Sea |
| Pouting | *Trisopterus luscus* | Froese and Pauly, 2017 | Froese and Pauly, 2017 | Merayo, 1998 | Merayo, 1998 | Merayo, 1998 | Barnes et al, 2008 | Mean FAO (GFCM) and Sea Around Us | | Other |
| Poor cod | *Trisopterus minutus* | Froese and Pauly, 2017 | Froese and Pauly, 2017 | Merayo, 1998 | Tsikliras et al, 2010 | Merayo, 1998 | Barnes et al, 2008 | Mean FAO (GFCM) and Sea Around Us | | Med Sea |
| Goldband goatfish | *Upeneus moluccensis* | Froese and Pauly, 2017 | Tsikliras and Stergiou, 2013 | Saad, 1998 | Tsikliras et al, 2010 | By default | Barnes et al, 2008 | Mean FAO (GFCM) and Sea Around Us | | Other |
| Swordfish | *Xiphias gladius* | ICCAT 2016 | ICCAT 2016 | Macias et al, 2005 | Tsikliras et al, 2010 | Macias et al, 2005 | Barnes et al, 2008 | ICCAT 2016 | ICCAT 2016 | Med Sea |
| Grass goby | *Zosterisessor ophiocephalus* | Froese and Pauly, 2017 | Tsikliras and Stergiou, 2013 | Franco et al, 2002 | Tsikliras et al, 2010 | By default | Barnes et al, 2008 | Mean FAO (GFCM) and Sea Around Us | | Med Sea |

### Appendix D. Modelling high trophic level species distribution

Table D.2. Species name, number of used occurrence compiled from the OBIS and GBIF database using the "spocc" R package (https://github.com/ropensci/spocc) and the robis R package (https://github.com/iobis/robis). The average True Skill Statistic (TSS) values from the ensemble modelling approach is given. According to the Landis and Koch (1977) accuracy classification scheme, TSS can be classified as follows: excellent, TSS>0.8; good, 0.6<TSS<0.8; fair, 0.4<TSS<0.6; poor, 0.2<TSS<0.4; and no predictive ability, TSS<0.2 (Ben Rais Lasram et al., 2010).

| Species | Number of OBIS/GBIF records | Mean TSS of the ensemble modelling approach |
|---|---|---|
| Alosa_alosa | 490 | 0.689 |
| Alosa_fallax | 1014 | 0.994 |
| Anguilla_anguilla | 11902 | 0.782 |
| Argyrosomus_regius | 122 | 0.788 |
| Aristaeomorpha_foliacea | 783 | 0.95 |
| Aristeus_antennatus | 345 | 0.879 |
| Atherina_boyeri | 381 | 0.802 |
| Auxis_rochei_rochei | 473 | 0.823 |
| Belone_belone | 744 | 0.811 |
| Boops_boops | 3302 | 0.832 |
| Caranx_crysos | 14902 | 0.676 |
| Chelidonichthys_lucerna | 5580 | 0.823 |
| Coris_julis | 1291 | 0.764 |
| Coryphaena_hippurus | 48490 | 0.668 |
| Crangon_crangon | 8734 | 0.869 |
| Crystallogobius_linearis | 223 | 0.802 |
| Dentex_dentex | 175 | 0.857 |
| Dentex_gibbosus | 499 | 0.861 |
| Dentex_maroccanus | 194 | 0.834 |
| Dicentrarchus_labrax | 1685 | 0.864 |
| Diplodus_annularis | 1113 | 0.734 |
| Diplodus_cervinus | 254 | 0.659 |
| Diplodus_puntazzo | 238 | 0.759 |
| Diplodus_sargus_sargus | 676 | 0.82 |
| Diplodus_vulgaris | 1035 | 0.645 |
| Eledone_cirrhosa | 2009 | 0.856 |
| Engraulis_encrasicolus | 3649 | 0.876 |
| Epinephelus_aeneus | 746 | 0.793 |
| Epinephelus_marginatus | 1180 | 0.708 |
| Etrumeus_teres | 2935 | 0.71 |
| Eutrigla_gurnardus | 22987 | 0.862 |
| Galeus_melastomus | 1400 | 0.762 |
| Gobius_niger | 3286 | 0.69 |
| Halobatrachus_didactylus | 190 | 0.674 |
| Illex_coindetii | 192 | 0.851 |
| Lepidorhombus_whiffiagonis | 6340 | 0.879 |
| Liza_aurata | 257 | 0.998 |
| Liza_ramada | 424 | 0.833 |
| Liza_saliens | 77 | 0.799 |
| Loligo_vulgaris | 6471 | 0.903 |
| Lophius_budegassa | 2223 | 0.873 |
| Lophius_piscatorius | 9817 | 0.874 |
| Merlangius_merlangus | 34994 | 0.857 |
| Merluccius_merluccius | 11243 | 0.778 |
| Micromesistius_poutassou | 5791 | 0.667 |
| Mugil_cephalus | 24628 | 0.739 |
| Mullus_barbatus_barbatus | 999 | 0.876 |
| Mullus_surmuletus | 5104 | 0.882 |
| Mustelus_mustelus | 2459 | 0.88 |
| Nephrops_norvegicus | 4344 | 0.816 |
| Octopus_vulgaris | 2505 | 0.804 |
| Pagellus_acarne | 998 | 0.656 |
| Pagellus_erythrinus | 1676 | 0.791 |
| Pagrus_pagrus | 13517 | 0.64 |
| Palaemon_serratus | 3352 | 0.991 |
| Palinurus_elephas | 903 | 0.968 |
| Parapenaeus_longirostris | 1847 | 0.807 |
| Penaeus_kerathurus | 547 | 0.825 |
| Phycis_phycis | 733 | 0.626 |
| Platichthys_flesus_flesus | 18621 | 0.804 |
| Pleuronectes_platessa | 33110 | 0.869 |
| Pomatomus_saltatrix | 10309 | 0.879 |
| Pomatoschistus_marmoratus | 52 | 0.763 |
| Pomatoschistus_minutus | 5421 | 0.761 |
| Rhinobatos_rhinobatos | 242 | 0.886 |

| | | |
|---|---|---|
| Sarda_sarda | 1479 | 0.657 |
| Sardina_pilchardus | 4409 | 0.701 |
| Sardinella_aurita | 14094 | 0.87 |
| Saurida_undosquamis | 3521 | 0.788 |
| Sciaena_umbra | 304 | 0.805 |
| Scomber_japonicus | 6230 | 0.691 |
| Scomber_scombrus | 34671 | 0.872 |
| Scophthalmus_maximus | 7499 | 0.816 |
| Scorpaena_notata | 859 | 0.677 |
| Scyliorhinus_canicula | 17348 | 0.793 |
| Sepia_officinalis | 4292 | 0.81 |
| Seriola_dumerili | 14390 | 0.988 |
| Serranus_atricauda | 706 | 0.696 |
| Solea_solea | 11592 | 0.895 |
| Sparus_aurata | 922 | 0.85 |
| Sphyraena_sphyraena | 489 | 0.619 |
| Sphyraena_viridensis | 256 | 0.685 |
| Spicara_maena | 663 | 0.805 |
| Spicara_smaris | 908 | 0.999 |
| Spondyliosoma_cantharus | 1162 | 0.762 |
| Sprattus_sprattus | 21210 | 0.764 |
| Squilla_mantis | 330 | 0.981 |
| Stephanolepis_diaspros | 32 | 0.881 |
| Thunnus_alalunga | 23142 | 0.66 |
| Thunnus_thynnus | 7316 | 0.64 |
| Trachurus_mediterraneus | 1082 | 0.861 |
| Trachurus_picturatus | 2363 | 0.884 |
| Trachurus_trachurus | 19826 | 0.655 |
| Trachyrincus_scabrus | 350 | 0.824 |
| Trigla_lyra | 1054 | 0.831 |
| Trisopterus_luscus | 8650 | 0.806 |
| Trisopterus_minutus | 18418 | 0.776 |
| Upeneus_moluccensis | 788 | 0.764 |
| Xiphias_gladius | 82441 | 0.642 |
| Zosterisessor_ophiocephalus | 24 | 0.639 |

Figure D.1. Species distribution maps of OSMOSE-MED species built with a niche modelling approach using the BIOMOD2 R package (Thuillier et al., 2009). Presence/absence distribution were built by conserving the occurrence probabilities for pixels above the sensitivity-specificity sum maximization threshold (i.e. threshold that maximized the True Skill Statistic criterion).

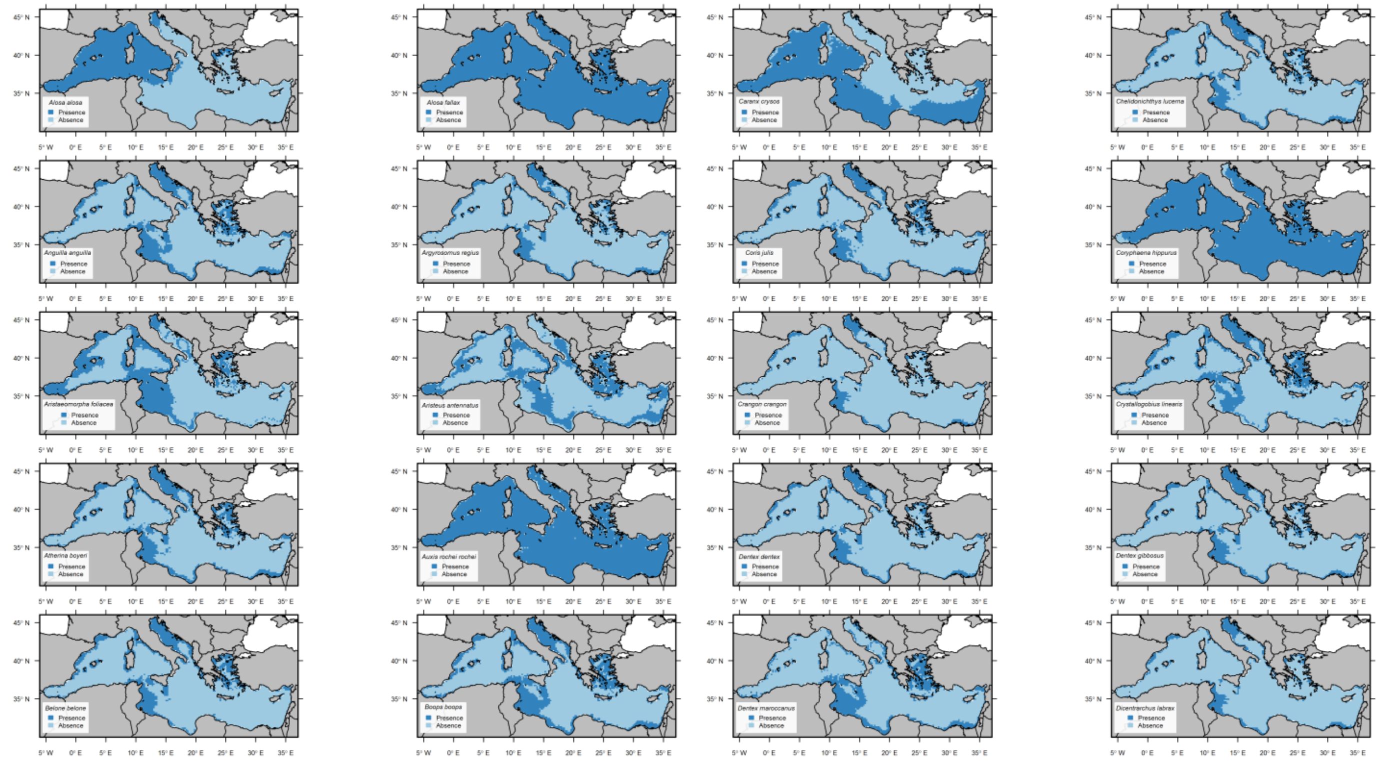

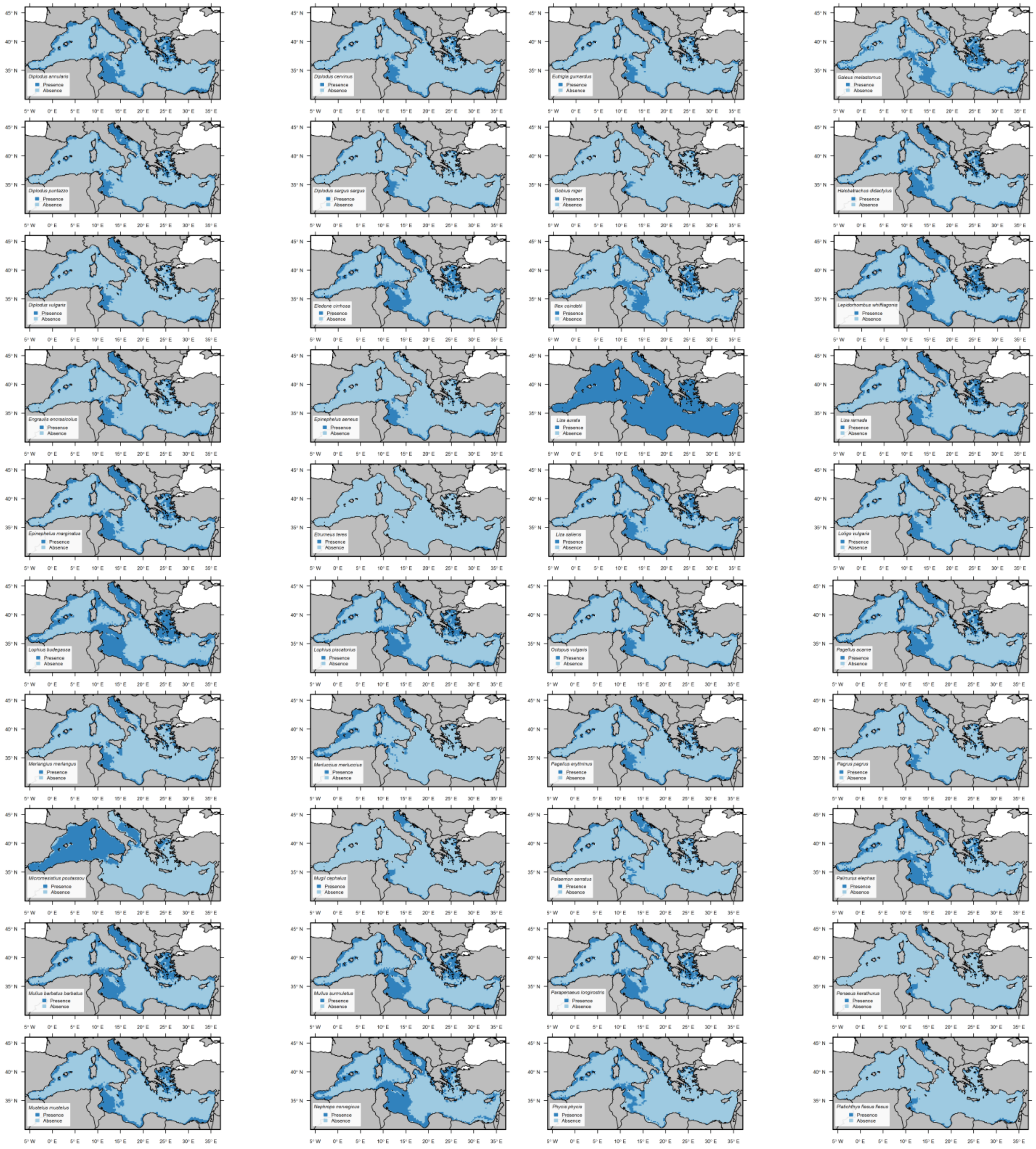
Diplodus annularis
Diplodus cervinus
Eutrigla gurnardus
Galeus melastomus
Diplodus puntazzo
Diplodus sargus sargus
Gobius niger
Halobatrachus didactylus
Diplodus vulgaris
Eledone cirrhosa
Illex coindetii
Lepidorhombus whiffiagonis
Engraulis encrasicolus
Epinephelus aeneus
Liza aurata
Liza ramada
Epinephelus marginatus
Etrumeus teres
Liza saliens
Loligo vulgaris
Lophius budegassa
Lophius piscatorius
Octopus vulgaris
Pagellus acarne
Merlangius merlangus
Merluccius merluccius
Pagellus erythrinus
Pagrus pagrus
Micromesistius poutassou
Mugil cephalus
Palaemon serratus
Palinurus elephas
Mullus barbatus barbatus
Mullus surmuletus
Parapenaeus longirostris
Penaeus kerathurus
Mustelus mustelus
Nephrops norvegicus
Phycis phycis
Platichthys flesus flesus
Presence
Absence
45° N
40° N
35° N
5° W
0° E
5° E
10° E
15° E
20° E
25° E
30° E
35° E

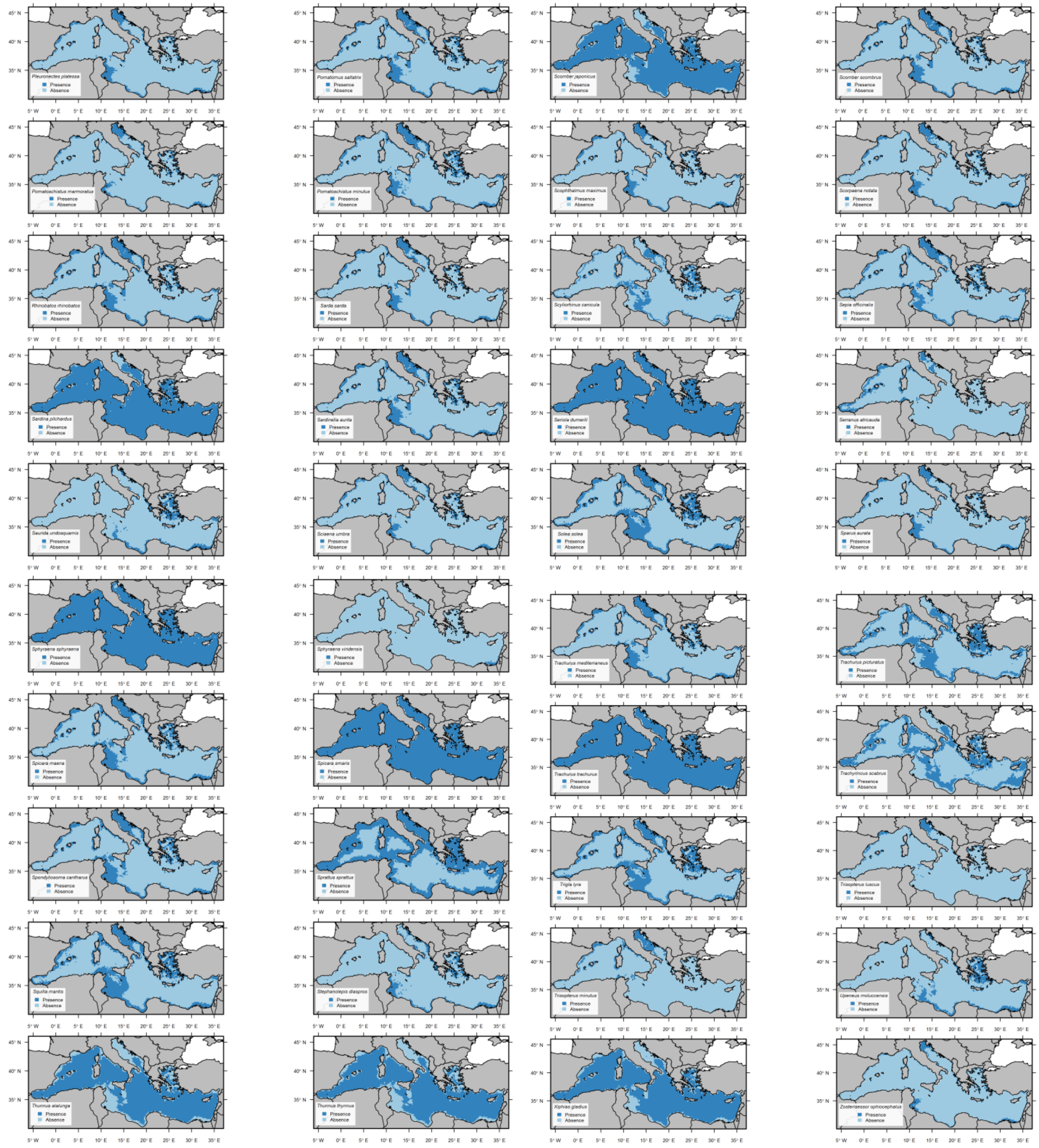

### Appendix E. Estimated parameters during the calibration of OSMOSE-MED

Table E.1. Availability coefficients of low trophic levels organisms (plankton and benthos) to high trophic levels organisms, fishing mortality and larval mortality rate parameters resulting form the calibration of OSMOSE-MED.

| Common name | Scientific name | coefficient of plankton accessibility | Fishing mortality (*year*$^{-1}$) | Larval mortality (*year*$^{-1}$) |
|---|---|---|---|---|
| Picophytoplankton | Synechococcus spp. | 0.11 | -- | -- |
| Nanophytoplankton | Dinoflagellates | $5.44 .10^{-7}$ | -- | -- |
| Microphytoplankton | Diatoms | $3.65 .10^{-6}$ | -- | -- |
| Nanozooplankton | Bacterivorous flagellates and small ciliates | $1.80 .10^{-8}$ | -- | -- |
| Microzooplankton | Ciliates and large flagellates | $2.64 .10^{-5}$ | -- | -- |
| Mesozooplankton | Copepods and amphipods | 0.099 | -- | -- |
| Benthos | -- | 0.055 | -- | -- |
| Allis shad | *Alosa alosa* | -- | 0.46 | 0.38 |
| Twaite shad | *Alosa fallax* | -- | 0.59 | 4.53 |
| European eel | *Anguilla anguilla* | -- | 1.13 | 9.02 |
| Meagre | *Argyrosomus regius* | -- | 0.58 | 4.17 |
| Giant red shrimp | *Aristaeomorpha foliacea* | -- | 0.06 | 0.21 |
| Blue and red shrimp | *Aristeus antennatus* | -- | 0.68 | 0.97 |
| Big-scale sand smelt | *Atherina boyeri* | -- | 0.03 | 0.56 |
| Bullet tuna | *Auxis rochei rochei* | -- | 1.67 | 8.66 |
| Garfish | *Belone belone* | -- | 1.17 | 2.50 |
| Bogue | *Boops boops* | -- | 0.64 | 1.58 |
| Blue runner | *Caranx crysos* | -- | 0.00 | 5.45 |
| Tub gurnard | *Chelidonichthys lucerna* | -- | 0.18 | 8.22 |
| Mediterranean rainbow wrasse | *Coris julis* | -- | 0.15 | 0.60 |
| Common dolphin fish | *Coryphaena hippurus* | -- | 2.42 | 8.81 |
| Common shrimp | *Crangon crangon* | -- | 0.13 | 0.44 |
| Cristal goby | *Crystallogobius linearis* | -- | 0.00 | 0.19 |
| Common dentex | *Dentex dentex* | -- | 1.01 | 3.31 |
| Pink dentex | *Dentex gibbosus* | -- | 0.35 | 6.44 |
| Morocco dentex | *Dentex maroccanus* | -- | 0.00 | 5.09 |
| European seabass | *Dicentrarchus labrax* | -- | 0.81 | 5.87 |
| Annular sea bream | *Diplodus annularis* | -- | 0.53 | 1.68 |
| Zebra seabream | *Diplodus cervinus* | -- | 0.34 | 3.21 |
| Sharpsnout seabream | *Diplodus puntazzo* | -- | 0.28 | 3.76 |
| White sea bream | *Diplodus sargus sargus* | -- | 0.38 | 1.21 |
| Common two-banded sea bream | *Diplodus vulgaris* | -- | 0.32 | 0.96 |
| Horned octopus | *Eledone cirrhosa* | -- | 0.56 | 3.09 |
| European anchovy | *Engraulis encrasicolus* | -- | 0.15 | 8.70 |
| White grouper | *Epinephelus aeneus* | -- | 0.34 | 9.71 |
| Dusky grouper | *Epinephelus marginatus* | -- | 0.79 | 1.50 |
| Red-eye round herring | *Etrumeus teres* | -- | 0.13 | 2.54 |
| Grey gurnard | *Eutrigla gurnardus* | -- | 0.11 | 0.57 |
| Black-mouthed dogfish | *Galeus melastomus* | -- | 0.40 | 2.24 |
| Black goby | *Gobius niger* | -- | 0.34 | 2.48 |
| Lusitanian toadfish | *Halobatrachus didactylus* | -- | 0.36 | 4.03 |
| Shortfin squid | *Illex coindetii* | -- | 0.51 | 8.70 |
| Megrim | *Lepidorhombus whiffiagonis* | -- | 0.16 | 3.30 |
| Golden grey mullet | *Liza aurata* | -- | 0.22 | 5.98 |
| Thinlip grey mullet | *Liza ramada* | -- | 0.00 | 4.51 |
| Leaping mullet | *Liza saliens* | -- | 0.00 | 2.36 |
| European squid | *Loligo vulgaris* | -- | 0.28 | 2.47 |
| Black-bellied angler | *Lophius budegassa* | -- | 0.20 | 5.80 |
| Anglerfish | *Lophius piscatorius* | -- | 0.77 | 2.53 |
| Whiting | *Merlangius merlangus* | -- | 0.41 | 1.60 |
| European hake | *Merluccius merluccius* | -- | 0.39 | 8.62 |
| Blue whiting | *Micromesistius poutassou* | -- | 1.43 | 2.12 |
| Flathead grey mullet | *Mugil cephalus* | -- | 1.57 | 3.13 |
| Red mullet | *Mullus barbatus barbatus* | -- | 0.97 | 1.14 |
| Striped red mullet | *Mullus surmuletus* | -- | 0.46 | 3.38 |
| Smooth hound | *Mustelus mustelus* | -- | 0.74 | 0.84 |
| Norway lobster | *Nephrops norvegicus* | -- | 0.28 | 0.41 |
| Common octopus | *Octopus vulgaris* | -- | 0.91 | 5.00 |
| Axillary seabream | *Pagellus acarne* | -- | 0.79 | 4.51 |
| Common pandora | *Pagellus erythrinus* | -- | 0.61 | 1.04 |
| Common seabream | *Pagrus pagrus* | -- | 0.37 | 0.72 |

| | | | | |
|---|---|---|---|---|
| Common prawn | *Palaemon serratus* | -- | 0.40 | 1.40 |
| Common spiny lobster | *Palinurus elephas* | -- | 0.20 | 4.72 |
| Deep-water rose shrimp | *Parapenaeus longirostris* | -- | 0.29 | 2.01 |
| Caramote prawn | *Penaeus kerathurus* | -- | 0.75 | 0.14 |
| Forkbeard | *Phycis phycis* | -- | 0.61 | 3.57 |
| European flounder | *Platichthys flesus flesus* | -- | 0.67 | 6.83 |
| European plaice | *Pleuronectes platessa* | -- | 0.00 | 6.13 |
| Bluefish | *Pomatomus saltatrix* | -- | 0.84 | 4.67 |
| Marbled goby | *Pomatoschistus marmoratus* | -- | 0.00 | 0.51 |
| Sand goby | *Pomatoschistus minutus* | -- | 0.00 | 0.30 |
| Common guitarfish | *Rhinobatos rhinobatos* | -- | 0.23 | 7.85 |
| Atlantic bonito | *Sarda sarda* | -- | 0.90 | 8.42 |
| European pilchard | *Sardina pilchardus* | -- | 0.49 | 3.32 |
| Round sardinella | *Sardinella aurita* | -- | 1.36 | 3.66 |
| Brushtooth lizardfish | *Saurida undosquamis* | -- | 0.14 | 4.25 |
| Brown meagre | *Sciaena umbra* | -- | 0.16 | 3.78 |
| Chub mackerel | *Scomber japonicus* | -- | 1.00 | 6.85 |
| Atlantic mackerel | *Scomber scombrus* | -- | 1.01 | 3.47 |
| Turbot | *Scophthalmus maximus* | -- | 0.74 | 5.20 |
| Small red scorpionfish | *Scorpaena notata* | -- | 0.00 | 0.74 |
| Lesser spotted dogfish | *Scyliorhinus canicula* | -- | 0.17 | 10.60 |
| Common cuttlefish | *Sepia officinalis* | -- | 2.06 | 5.54 |
| Greater amberjack | *Seriola dumerili* | -- | 2.09 | 3.72 |
| Blacktail comber | *Serranus atricauda* | -- | 0.00 | 6.27 |
| Common sole | *Solea solea* | -- | 0.63 | 6.48 |
| Gilthead seabream | *Sparus aurata* | -- | 1.04 | 0.56 |
| European barracuda | *Sphyraena sphyraena* | -- | 1.10 | 2.78 |
| Yellowmouth barracuda | *Sphyraena viridensis* | -- | 0.00 | 0.35 |
| Blotched picarel | *Spicara maena* | -- | 0.15 | 0.23 |
| Picarel | *Spicara smaris* | -- | 0.59 | 1.13 |
| Black seabream | *Spondyliosoma cantharus* | -- | 1.34 | 2.62 |
| European sprat | *Sprattus sprattus* | -- | 0.21 | 1.24 |
| Spottail mantis shrimp | *Squilla mantis* | -- | 0.23 | 3.73 |
| Reticulated leatherjacket | *Stephanolepis diaspros* | -- | 0.00 | 3.61 |
| Albacore | *Thunnus alalunga* | -- | 0.09 | 4.30 |
| Atlantic bluefin tuna | *Thunnus thynnus* | -- | 0.07 | 2.22 |
| Mediterranean horse mackerel | *Trachurus mediterraneus* | -- | 0.80 | 0.88 |
| Blue jack mackerel | *Trachurus picturatus* | -- | 0.54 | 2.66 |
| Atlantic horse mackerel | *Trachurus trachurus* | -- | 1.13 | 3.73 |
| Piper gurnard | *Trigla lyra* | -- | 0.02 | 6.68 |
| Pouting | *Trisopterus luscus* | -- | 0.60 | 1.68 |
| Poor cod | *Trisopterus minutus* | -- | 0.60 | 0.84 |
| Goldband goatfish | *Upeneus moluccensis* | -- | 0.07 | 7.52 |
| Swordfish | *Xiphias gladius* | -- | 0.54 | 6.68 |
| Grass goby | *Zosterisessor ophiocephalus* | -- | 0.00 | 2.97 |
| Euphausiids | - | -- | 0.00 | 2.63 |

### Appendix F. Details on the MEDITS demersal survey

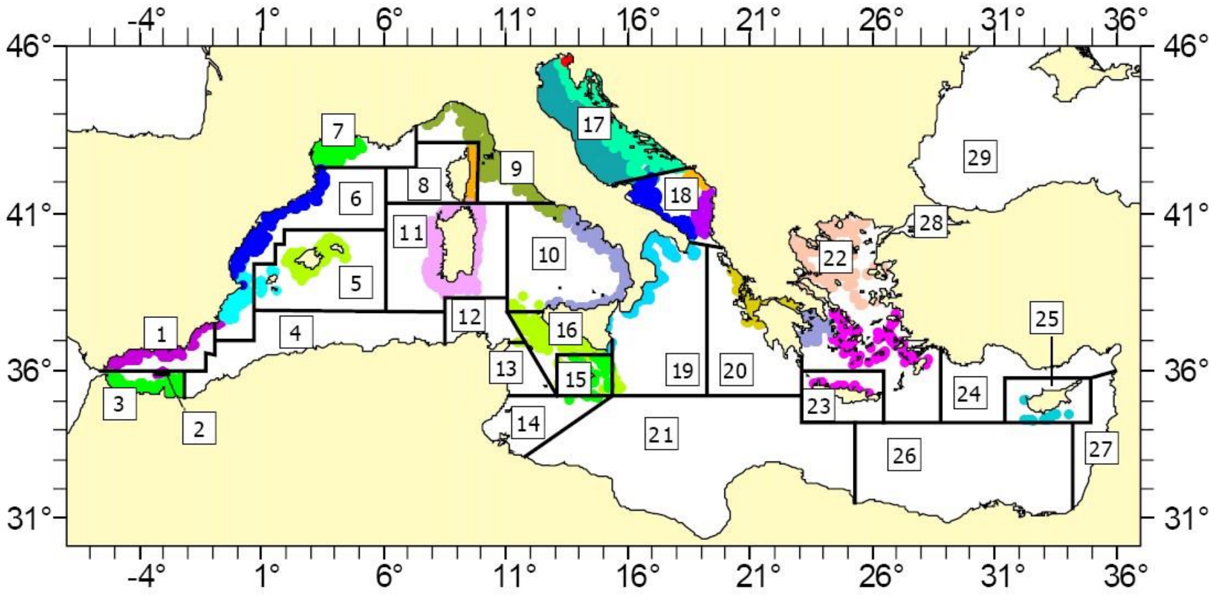


*Figure F.2. Study area and sampling sites of trawl surveys based on the MEDITS program protocol. From* [*http://www.sibm.it/SITO%20MEDITS/principalemedits.htm*](http://www.sibm.it/SITO%20MEDITS/principalemedits.htm)

Table F.1. Details on Geographical Sub-Areas (GSAs) for which biomass estimates from MEDITS survey were available on the 2006-2013 period. MEDITS survey data can be available on request to national coordinators of the program.

| GSA numbers | GSA names | GFCM subregions | GSA surfaces (km²) |
|---|---|---|---|
| 1-2 | Northern Alboran Sea and Alboran island | Western Mediterranean Sea | 35267 |
| 5 | Balearic island | | 113921 |
| 6 | Northern Spain | | 101577 |
| 7 | Gulf of Lions | | 62669 |
| 8 | Corsica island | | 25736 |
| 9 | Ligurian and North Tyrrhenian Sea | | 42410 |
| 10 | South Tyrrhenian Sea | | 134365 |
| 11 | West and East Sardinia | | 125731 |
| 18 | Southern Adriatic Sea | Adriatic Sea | 52610 |
| 16 | Southern Sicily | Ionian Sea | 37431 |
| 19 | Western Ionian Sea | | 167046 |
| 20 | Eastern Ionian Sea | | 124279 |
| 22 | Aegean Sea | Eastern Mediterranean Sea | 223337 |
| 23 | Crete island | | 62680 |
| 25 | Cyprus island | | 45100 |
| | | Total surface of available GSAs | 1366001 |
| | | % of the surface of the OSMOSE-MED model | 55 % |
